# Controlled Fluorescent Labelling of Metal Oxide Nanoparticles for Artefact-free Live Cell Microscopy

**DOI:** 10.1101/2021.04.19.440400

**Authors:** Boštjan Kokot, Hana Kokot, Polona Umek, Katarina Petra van Midden, Stane Pajk, Maja Garvas, Christian Eggeling, Tilen Koklič, Iztok Urbančič, Janez Štrancar

## Abstract

Nanotechnologies hold great promise for various applications. To predict and guarantee the safety of novel nanomaterials, it is essential to understand their mechanism of action in an organism, causally connecting adverse outcomes with early molecular events. They are best investigated using non-invasive advanced optical methods, such as high-resolution live-cell fluorescence microscopy, which require stable labelling of nanoparticles with fluorescent dyes. When performed inadequately, unbound fluorophores and inadvertently altered chemical and physical properties of the nanoparticles can, however, result in experimental artefacts and erroneous conclusions.

To prevent such unintentional errors, we here describe a minimal combination of experimental methods to enable artefact-free fluorescent labelling of metal-oxide nanoparticles – the largest subpopulation of nanoparticles by industrial production and applications – and demonstrate its application in the case of TiO_2_ nanotubes. We 1) characterize potential changes of the nanoparticles’ surface charge and morphology that might occur during labelling, and 2) assess stable binding of the fluorescent dye to nanomaterial, which ensures correct nanoparticle localization. Together, these steps warrant the reliability and reproducibility of advanced optical tracking, which is necessary to explore nanomaterials’ mechanism of action and will foster widespread and safe use of new nanomaterials.

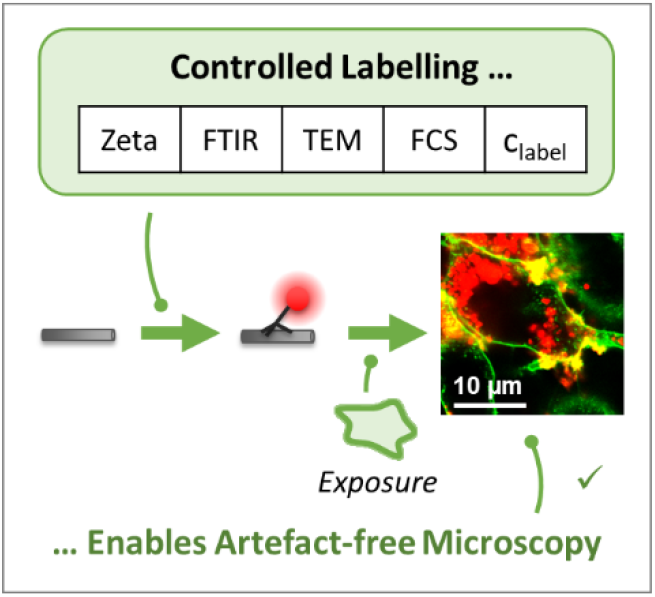

## 1. Introduction

The rapid development of new materials is inevitably accompanied by their processing, combustion, and degradation, which can release particulate matter into the environment and increase our daily exposure to nanomaterials. ^[1–3]^ Moreover, nanomaterials are becoming ever more widely used in various industrial branches, including pharmacy ^[4,5]^ and the food industry ^[6,7]^ – all due to their unique properties. Consequently, there is a growing concern regarding the safety of nanomaterials, urging mandatory testing for their potential toxic effects. ^[8,9]^ However, it has been recognized that many nanomaterials interfere with classic cytotoxicity essays, ^[10]^ troubling their safety assessments and hence hindering their development.

To ease the introduction of safe nanomaterials into our daily lives, the scientific community has recognized the need for a deeper understanding of the link between adverse outcomes (nanoparticle toxicity) and nanoparticle characteristics. ^[11–14]^ To unravel fundamental molecular mechanisms of nanoparticle toxicity, it is crucial to identify the molecular events and their causal relationships, which heavily relies on monitoring nanoparticle localization and interaction with living cells in real-time. ^[15–19]^

One of the most powerful experimental techniques to study early molecular events including nanoparticle localization and interaction with living cells is fluorescence microscopy, especially its advanced super-resolution implementation with spatial and temporal resolution on the order of tens of nanometers and seconds, respectively. However, since most of the nanoparticles are not intrinsically fluorescent, such an experiment requires special preparation of the nanomaterial. Here, we will focus on fluorescent labelling which binds fluorescent dyes to the nanoparticle. As most commercially available fluorescent dyes bind weakly onto the native surface of the majority of nanoparticles, a commonly-used approach is to functionalize the nanoparticle surface with a coupling agent to create a link between the inorganic surface of the nanoparticle and the organic fluorescent molecule. ^[20–22]^ An alternative approach, luminescence labelling, can be utilized by doping the nanomaterial with lanthanides during its synthesis, ^[23,24]^ which increases the photostability of the labelled nanomaterial and minimizes the lanthanide dissolution. However, the multistep synthesis of the doped nanomaterial is lengthy, requires optimization for every dopant separately, yields less bright fluorescence, and, unfortunately, cannot be applied to already synthesized nanomaterials.

In order for our method to be applicable to the widest range of nanomaterials, the labelling procedure that is assessed here, is assumed to consist of functionalization with a linker, attachment of the fluorescent dye, and removal of the free dye (**Figure 1a**). Noteworthy, though, each of these steps can inadvertently change the properties of the nanomaterial, such as its morphology, surface charge, and the quantity of non-specifically adhered dye, ^[25,26]^ thereby also altering the nanoparticle’s interactions with the environment and hence affecting the interpretation of a fluorescence localization and tracking experiment (Figure 1b). Thus, reproducibility and reliability of experiments with fluorescent nanomaterial can be assured only if the labelling procedure of nanoparticles is carefully controlled at each step that can potentially alter the nanomaterial properties.

**Figure 1.**
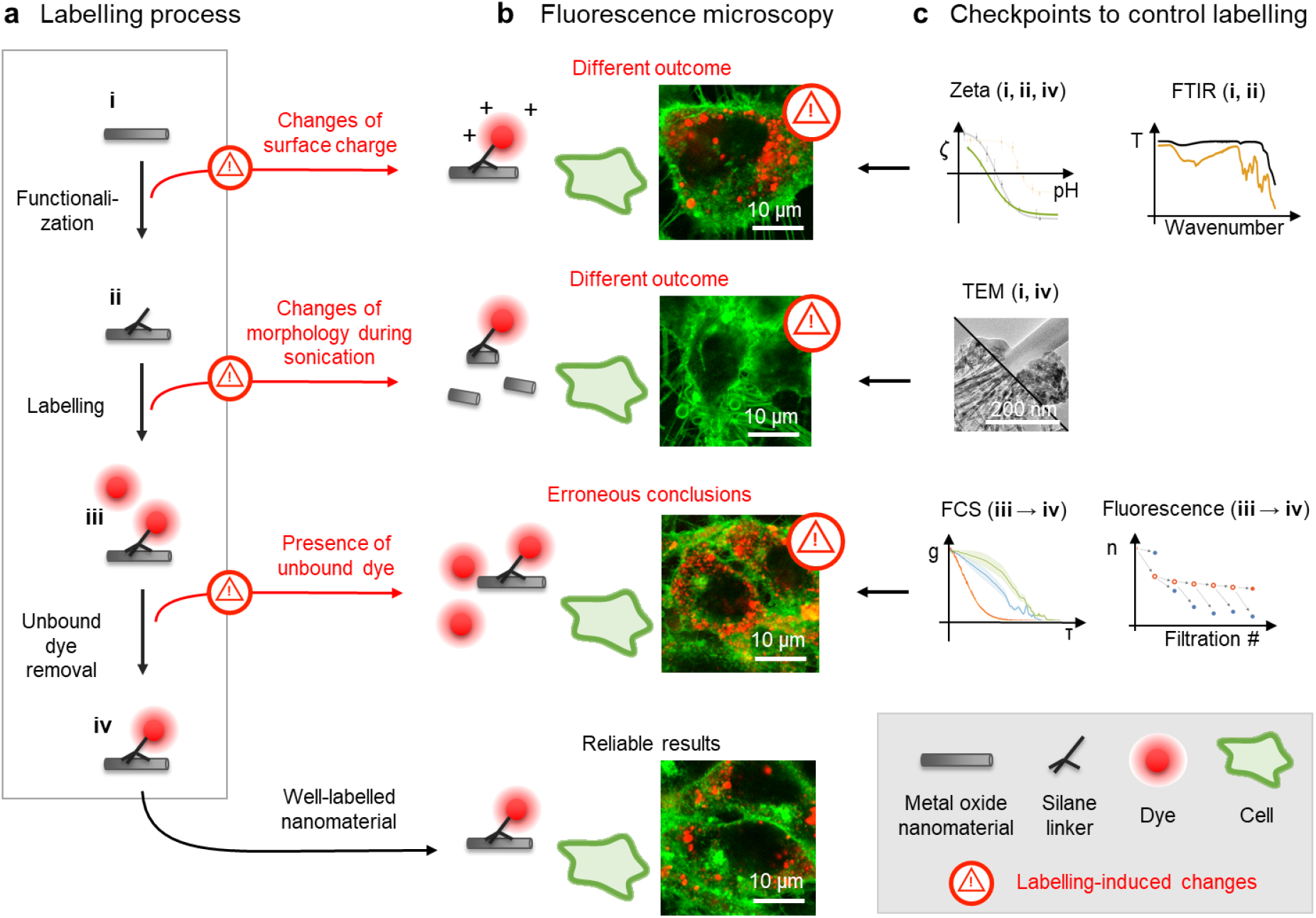
Overview of nanomaterial labelling with fluorescent dyes for live cell imaging, with checkpoints to prevent inadvertent changes to the original nanomaterial. **a** Schematic of nanomaterial labelling: the initial nanomaterial (i) is functionalized with a linker (ii), to which a fluorescent dye is bound (iii), and finally unbound dye is washed away (iv). Note that the scheme elements are not drawn to scale. **b** Comparison of fluorescence micrographs of LA-4 cells (membranes labelled with CellMask Orange, shown in green) incubated with the exemplary nanomaterial – TiO_2_ nanotubes – that were either adequately (bottom-most) or inadequately (top three images) fluorescently labelled with Alexa Fluor 647 (shown in red). Inadequate labelling resulting in different outcomes of the experiments and thus possibly leads to erroneous conclusions. For micrographs of separate color channels refer to Figure S4 in Supporting Information. **c** Proposed checkpoints to prevent such errors by measuring the nanomaterial properties at several stages during the labelling procedure.

Several cases of fluorescent labelling or doping of different nanoparticles, ranging from gold and metal oxides to carbonaceous and polystyrene nanoparticles, have been reported, with various degrees of quality control and nanomaterial characterization. To characterize the nanomaterial morphology, labelling protocols of nanoparticles are often paired with scanning-(SEM) and/or transmission-electron microscopy (TEM) of either the native surface before labelling ^[20,22,27,28]^ or of labelled nanoparticles. ^[29]^ When the dye is added during nanomaterial synthesis, often only the labelled nanomaterial is characterized. ^[25,26,30–35]^ Importantly, one must be aware that the labelling procedures often involve vigorous sonication to disperse the nanomaterial, which can alter the morphology of nanoparticles, ^[36]^ as shown in Figure 1.

Further surface modifications can arise from functionalization of the nanomaterial with a charged linker and fluorescent dye, which can alter the surface charge of the nanoparticles. This can be determined by Zeta potential measurements but is usually measured for either non-labelled or labelled nanoparticles. ^[20,30,35,37]^ However, only the comparison of the two ^[22]^ can give additional information on the success of the functionalization and labelling as well as possible labelling-induced modifications to the nanomaterial. Even more, the pH-dependence of the Zeta potential of any sample is rarely determined over the full pH range, yet only its full assessment ^[22,32]^ would allow predicting the surface charge of the nanoparticles in various cellular compartments with varying pH – for instance, in the phagolysosome; ^[38]^ such information can predictably impact the interpretation of experimental results. Besides Zeta potential, many other experimental methods can also test the success of the nanoparticle’ surface functionalization: Fourier transform infrared spectroscopy (FTIR), ^[22,39–43]^ high angle annular dark-field scanning transmission electron microscopy (HAADF-STEM), ^[22]^ secondary ion mass spectrometry (SIMS), ^[43]^ energy dispersive X-ray spectroscopy (EDX) ^[43]^ and/or X-ray photoelectron spectroscopy (XPS). ^[39,40]^ Although crucial, such a step is often omitted.

With time and/or change of the environment of the labelled nanoparticles, the linker or the fluorescent dye may desorb from the nanoparticles, potentially leading to experimental artefacts due to the unbound dye being indistinguishable from labelled nanomaterial on fluorescence images (Figure 1b). ^[25,26,35,44]^ Various methods are useful to remove the free fluorescent dye, including dialysis, ^[22,44]^ gel filtration, ^[25]^ centrifugal filtration ^[17,31,32]^ or a sequence of centrifugation and supernatant-removal. ^[20,33]^ The latter two methods enable precise characterization of free dye removal and the dye desorption from the nanoparticles by analyzing filtrates or supernatants, respectively. In the other approaches mentioned, dye desorption must be tested in an additional separate step. ^[26,35,44]^

To prevent erroneous conclusions from experiments performed with inadequately labelled and/or inadvertently changed nanomaterial, we propose and describe a minimal list of essential experimental checkpoints that ensure the labelling quality, characterize nanomaterial modification during labelling, and determine the rate of dye desorption (Figure 1c). These checkpoints should be measured at all major stages of the labelling procedure:

i. for native nanomaterial (Zeta potential, FTIR and TEM),
ii. for functionalized nanomaterial (Zeta potential and FTIR),
iii. → **iv** during free dye removal (FCS and fluorescence intensity), and
iv. for labelled nanomaterial (TEM and Zeta potential)

(see also the schematic in Figure S1 and flowchart in Figure S3 in Supporting Information). We demonstrate their application in the case of fluorescent labelling of TiO_2_ nanotubes, highlighting possible consequences of skipping the checkpoints at each step. For reference, the step-by-step labelling protocol is included in Supporting Information.

## 2. Results and Discussion

### 2.1. Characterization of Pristine Material

Before the labelling process, pristine nanomaterial (**i**) must first be thoroughly characterized. Even if provided by the manufacturer, it is recommended to measure the nanoparticle’s specific surface, elemental composition, and structure, for example by using the BET method (Brunauer–Emmett–Teller method), EDXS (Energy-dispersive X-ray spectroscopy), and XRD (X-ray diffraction), respectively. It is also important to be aware of the method of nanomaterial synthesis and chemicals that were used during the process because these chemicals can incorporate into or adsorb to the nanomaterial, contributing to its properties. For our TiO_2_ nanotubes, this has extensively been done and described previously. ^[22]^

At this stage, it is also crucial to determine the morphology of the native material with TEM (transmission electron microscopy) and its surface properties by measuring the pH dependence of the Zeta potential over the entire pH range. As will be discussed later on, these results are compared with the material’s properties during and after labelling to characterize potential labelling-induced modifications of morphology and surface charge.

### 2.2. Functionalization of Nanomaterial

Because fluorescent dyes do not bind covalently to the native surface of most metal oxides, the nanoparticles first need to be functionalized by binding a small linking chemical compound (linker) to the native nanoparticle surface (**Figure 2a, i** → **ii**). The amount of linker used should enable efficient labelling (i.e., the labelled nanomaterial has enough dyes attached to be detected in the experiments) while minimizing changes to the native surface of nanomaterial. The step-by-step protocol for the functionalization of TiO_2_ nanotubes with 3-(2-aminoethylamino)propyltrimethoxysilane (AEAPMS) is provided in the Supporting Information, whereas the procedures for some other nanoparticles and linkers can be found elsewhere. ^[39,39,42,43,45–47]^

**Figure 2.**
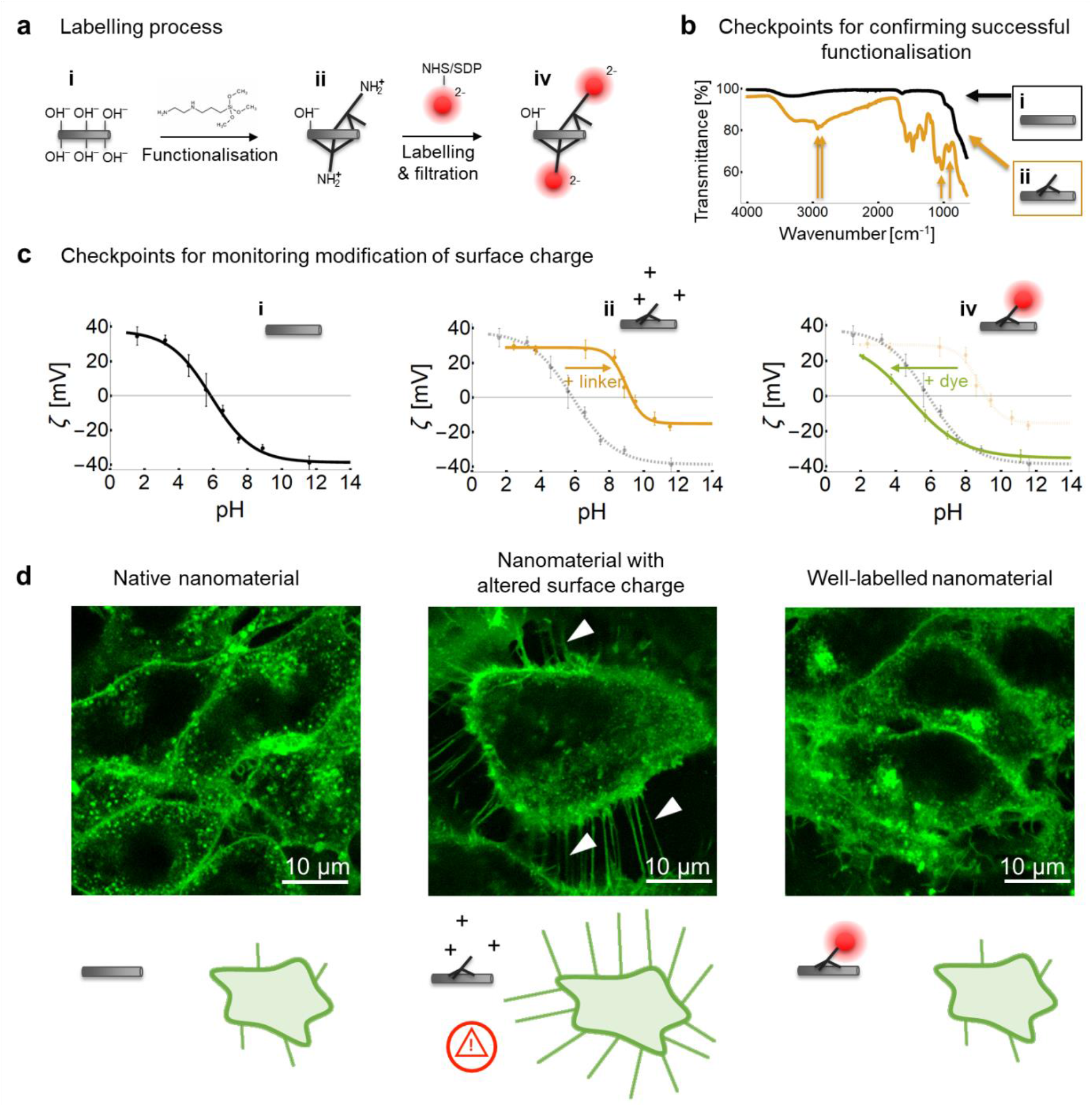
Functionalization and labelling of TiO_2_ nanoparticles with AEAPMS and Alexa Fluor dyes. **a** The schematics shows the chemical reactions taking place on the surface of nanoparticles upon their functionalization and subsequent fluorescent labelling, highlighting the dissociating chemical groups and their charges. **b** An ATR-FTIR spectrum of pristine (shown in black, **i**) and successfully functionalized TiO_2_ nanoparticles (shown in brown, **ii**), with corresponding schematics of the samples. Brown arrows indicate vibration bands of the chemical bonds of the AEAPMS linker attached to the TiO_2_ surface. Reference spectra of AEAPMS and solvents are shown in Figure S6, Supporting Information. **c** The pH-dependence of the Zeta potential (ζ) of **i** native TiO_2_ nanotubes, **ii** functionalized nanotubes, and **iv** nanotubes labelled with Alexa Fluor 488. Arrows above the graphs indicate the shift in Zeta potential between different stages. **d** Comparison of confocal fluorescence micrograph images of LA-4 epithelial cells (cell membranes are labelled with CellMask Orange and shown in green) after a 2-day exposure to (from left to right): unlabeled TiO_2_ nanotubes, functionalized nanotubes with a more positive surface charge, and TiO_2_ nanotubes labelled with Alexa Fluor 647 (its fluorescence is not shown), whose surface resembles the native nanomaterial. The ratio between the surface of the nanomaterial and surface of cells (so-called surface dose) was 10:1, 1:1, and 1:1, respectively. Notice the long filaments on the cell in the middle micrograph (arrowheads) which occur when the surface charge of the nanomaterial is changed but are absent in the micrographs of both the native and well-labelled nanomaterial.

When using a silane linker, in our case AEAPMS, the functionalization must be done in water-free toluene to prevent the formation of a water layer on the nanoparticle surface, which would disable the interaction between the linker and OH groups on the nanoparticles’ surface due to charge screening, ^[43]^ resulting in unsuccessful functionalization. The functionalized nanomaterial should be dried and stored as a powder to prevent desorption of the linker during storage.

#### 2.2.1. Characterization of Nanomaterial Functionalization

The success of functionalization is assessed using attenuated total reflection Fourier-transform infrared spectroscopy (ATR-FTIR) of the native (**i**) and functionalized nanomaterial (**ii**), as shown in Figure 2b for the pristine (shown in black) and successfully functionalized TiO_2_ nanotubes (shown in brown). If the functionalization has been successful, the spectrum of the functionalized nanomaterial should contain additional peaks, corresponding to the chemical bonds created by the binding of the linker to the nanomaterial with respect to the sum of the spectra of the linker and pristine nanomaterial (see Figure 2b and Figure S6, Supporting Information). ^[22]^ For comparison, an ATR-FTIR spectrum of poorly functionalized TiO_2_ nanotubes using the same AEAPMS linker is shown in Figure S6, Supporting Information, containing only low intensity of characteristic peaks of the linker-to-nanomaterial bonds.

At this stage, the pH-dependence of the Zeta potential of the functionalized nanomaterial is measured again (Figure 2c, **ii**). By comparing these pH dependencies to those obtained for the native nanomaterial (Figure 2c, **i**), the effect of the linker to the nanomaterial surface charge is further analyzed. For example, successful functionalization with the AEAPMS linker, which contains amino groups, shifts the isoelectric point (value at which the Zeta potential equals 0 mV) towards higher pH values (Figure 2c **ii**, orange arrow). If none or only a small amount of the linker has attached to the nanomaterial, the Zeta potential would remain unchanged – as shown in Figure S7 in Supporting Information.

Although using both FTIR and Zeta potential measurements may seem redundant, each of them brings important clues into the assessment of the labelling. On one hand, FTIR provides the required specificity in detecting the bond between the linker and the nanomaterial, while distinguishing unbound linkers from solvents and impurities. On the other hand, Zeta potential measurements of dispersed nanomaterial provide crucial information for choosing the appropriate charge of the fluorescent dye which shall counterbalance the charge effect of the linker to restore the surface charge of the native nanomaterial.

#### 2.2.2. Potential Artefacts

Importantly, the amount and charge of both the linker and the dye determine the final surface charge of the labelled nanomaterial, majorly influencing its interactions and mechanism of action in the cells and our bodies. ^[48,49]^ An example of this is shown in Figure 2d for TiO_2_ nanotubes with different surface charges. The morphology of the lung epithelial cells after a 2-day incubation with positively charged nanotubes is remarkably different than the morphology observed when they were incubated with native or labelled nanomaterial, indicating an effect of modified surface charge on the cell response to the nanomaterials. However, if the nanotubes’ Zeta potential was not determined before the measurement, these morphological changes and other toxicological outcomes could be wrongly attributed to the native nanomaterial.

### 2.3. Fluorescent Labelling

#### 2.3.1. Selection of the Fluorescent Dye

After verifying the functionalization step, the nanomaterial is ready to be labelled with the desired fluorescent dye. The choice of the dye must adhere to several criteria, and may even affect the choice of the linker. In short, the dye should be capable of forming a covalent bond with the linker, be appropriately charged to restore the surface charge of the labelled material as close to native as possible, and have suitable characteristics for the planned optical experiments – all of which we discuss in detail below.

Regarding the binding chemistry, many commercial dyes are offered with various reactive groups: NHS and SDP esters react with amines, maleimides react with thiol groups, hydrazides react with aldehydes and ketones, and azides and alkynes are used in “click” reactions, to name a few.

The spectral characteristics of fluorophores must match the wavelengths of the excitation light, dichroic and emission filters of the fluorescence microscope on which the experiments are to be done (see example in Figure S10 in Supporting Information). Moreover, it is advised that their spectra are well separated from the spectra of other fluorophores that will be used simultaneously in the experiments to minimize cross-talk and bleed-through, which can give rise to false co-localization and misinterpretations if not appropriately corrected for. ^[50–52]^ In live-cell applications, the dye should be bright and photo-stable to minimize the necessary excitation light flux, which can interfere with cellular processes during the experiment. ^[53]^ For use with advanced optical techniques (such as high-resolution microscopies, two-photon microscopy, fluorescence lifetime imaging, spectral imaging or correlation spectroscopy), further photo-physical aspects (e.g., lifetime, blinking, and environmental sensitivity) must also be taken into account.

Importantly, it is also necessary to select a dye which – in its unbound state – interacts with the sample as weakly as possible. Careful control of the number of dye molecules per nanomaterial can further decrease the interaction between the dye and the biological system. In general, it is always advisable to test the interaction between the dye and the sample with a control measurement, where the system is incubated with free dye instead of labelled nanomaterial for the desired amount of time. ^[54]^ By comparing the distribution and fluorescence intensity of the free dye with that of labelled nanomaterial, one can estimate the degree of false nanoparticle localization in the experiment.

Applying all these criteria to labelling of TiO_2_ nanoparticles, the dye should be an NHS (N-Hydroxysuccinimide) or SDP (sulfodichlorophenol) ester for covalent binding to the amine NH_2_ group on the AEAPMS linker, and have a negative charge to compensate the positive charge of the linker. We further wanted the selected dye to be compatible with our high-resolution STED microscope, for which bright and photo-stable far-red dyes work best. Also, interaction of the free dye with the system of interest needs to be taken into account: the unbound selected dye Alexa Fluor 647 does not stain the cell membrane due to a low membrane interaction factor, ^[55]^ and only a negligible fraction of the dye was localized in the cell after a 2-day incubation (Figure S18 Supporting Information).

#### 2.3.2. Fluorescent Labelling of Nanomaterials

During the labelling step, when fluorescent dye is to be covalently bound to the linker on the nanomaterial (Figure 1, **ii** → **iii)**, special care must be taken to ensure appropriate conditions for maximal labelling efficiency. Functionalized TiO_2_ nanoparticles and excess of fluorescent dye are combined according to the guidelines of the dye manufacturer. E.g., dyes with NHS and SDP esters must be stored in a water-free environment before labelling. For long-term storage aliquot the freshly opened dye in DMF (N,N-dimethylformamide), remove the DMF solvent under high-vacuum (10^−5^ bar) or lyophilization, and store under an inert gas (e.g. argon) at -70 °C. Just prior to labelling we dissolved the lyophilized probe in anhydrous DMSO (dimethyl sulfoxide).

Additional nanomaterial-specific requirements need to be considered: 1) To avoid nanoparticle aggregation during labelling, the concentration of nanoparticles should be kept below 1 mg/mL, the mixture should be sonicated by tip sonicator while labelling, and the pH of the medium should be in the range where the nanoparticles are stable. The appropriate pH value can be deduced from the Zeta potential measurements from the previous step; 2) To improve the labelling efficiency, the linker and the fluorescent dye should be oppositely charged at the pH of the labelling medium, and the osmolarity of the labelling medium should be kept as low as possible to avoid charge screening.

#### 2.3.3. Removal of the Free and Desorbed Dye

Because the labelling is performed with an excess of fluorescent dye, the dye that did not bind to the nanomaterial must be removed before experiments to increase the contrast and avoid false localization of nanomaterial in fluorescence images. For the same reason, we also have to remove the dye that has desorbed from the fluorescently labelled nanomaterial during storage – typically due to desorption of the linker from the nanomaterial’s surface, not cleavage of the dye from the linker. ^[43]^ The free dye can be conveniently removed either with a sequence of centrifugations and supernatant removals, or using a centrifugal filter device that will let unbound dye pass through. Both of these methods are quicker and use lower volumes of the washing media than the commonly used dialysis, enabling more reliable monitoring of the free dye removal due to higher concentrations of the dye in the filtrates (more on that below).

The first few filtrations should be made in a solvent in which the dye is soluble and nanomaterial is still well dispersed (in our case, a mixture of ethanol and diluted bicarbonate buffer in a 70:30 mass ratio, to achieve pH of 10, and osmolarity equivalent to 100-times diluted bicarbonate buffer, see Materials and methods section for details). The last few filtrations should use a medium that is not toxic to cells and in which the nanomaterial can be stably stored for extended periods of time to remove the solvent (in our case, 100-times diluted bicarbonate buffer with pH 10). Also, note that in-between centrifugations, the dispersion of nanoparticles should be gently sonicated on an ultrasound bath to minimize their aggregation and to desorb them from the filter or container.

#### 2.3.4. Evaluation of Labelling and Free Dye Removal

The labelling efficiency, dye removal, and dye desorption from the nanomaterial can be evaluated either by fluorescence intensity measurements of the filtrates and retentates (remaining sample after filtration) or by fluorescence correlation spectroscopy (FCS) of the retentates, both of which we describe below. In the method using centrifugation and supernatant removal, supernatants correspond to filtrates and resuspended pellets to retentates. To this end, the FCS and fluorescence intensity measurements are more or less interchangeable; however, the equipment for fluorescence intensity measurements is more common than for FCS, which can, on the other hand, distinguish free dye from dye bound to nanoparticles in the same sample based on co-diffusion analysis. Hence, we usually use fluorescence intensity measurements during the filtration process and FCS for the final assessment of the labelled nanomaterial.

##### Evaluation by Fluorescence Intensity

The quantities of the dye in the filtrates and retentates after each filtration step are determined from the fluorescence intensities measurements, performed with a spectrophotometer, from which absolute amounts (i.e., in moles) are calculated, taking into account the measured sample volumes, intensity-concentration calibration curves, scattering of light on the nanomaterial, as well as the influence of different media on the detected signal (see Figure S13-S17 in Supporting Information). The amount of the bound dye in the intermediate steps can be estimated from the fluorescence signal of the filtered dye (blue symbols in **Figure 3a**) to minimize the amount of the fluorescent material used for monitoring. In the estimations we neglected losses of the dye due to its adsorption to the laboratory glassware and plastics, but nevertheless reached a good agreement with the actually measured values (in Figure 3a, empty orange symbols extrapolate well towards the solid symbol); for more sticky dyes, though, additional calibration and correction may be required.

**Figure 3.**
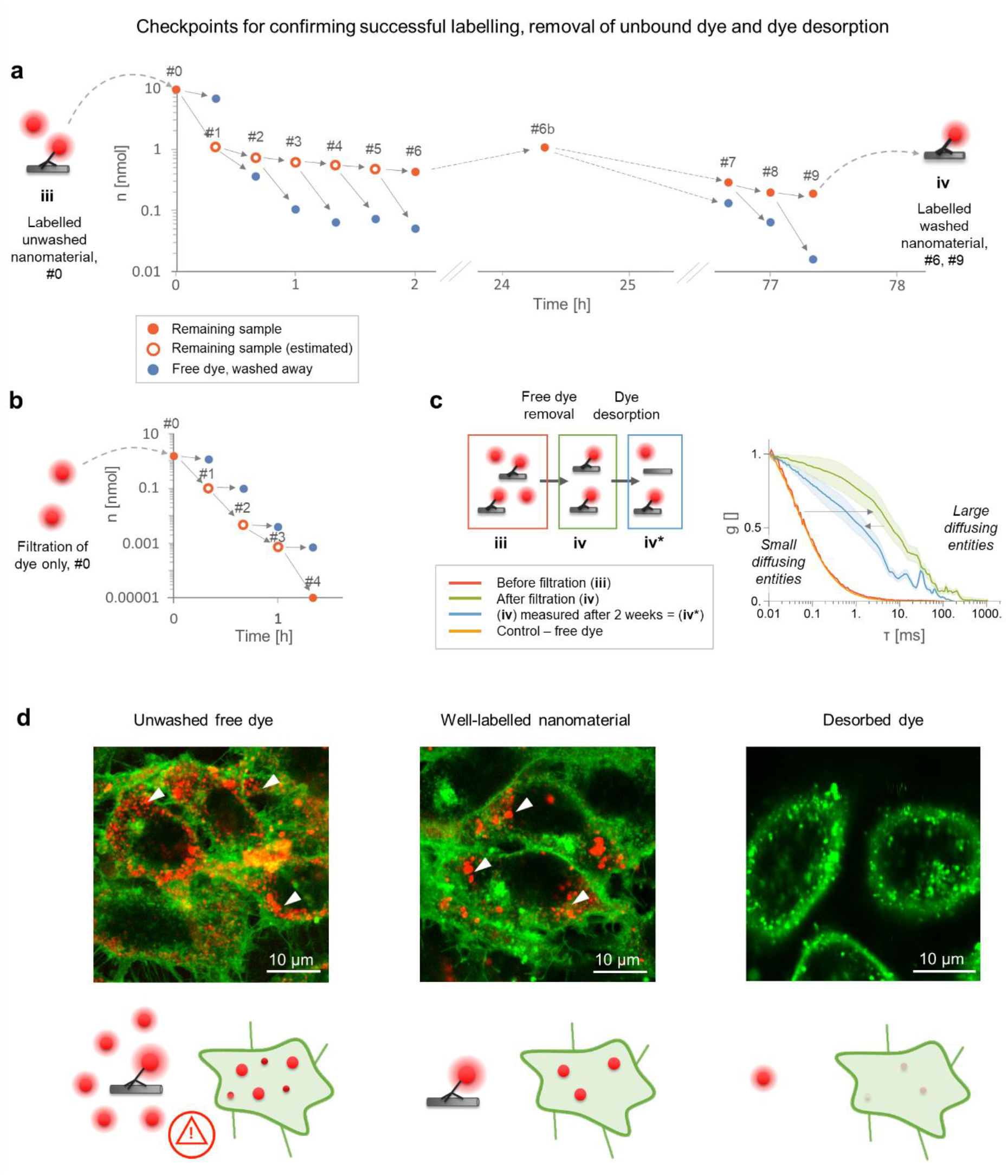
Removal of the free dye. **a** The strategy to control the removal of the free dye from the sample (TiO_2_ nanotubes, labelled with Alexa Fluor 488) by centrifugal filtration relies on monitoring the quantity (n [nmol]) of the washed-away dye (blue markers) as well as the quantity of the dye remaining in the sample (orange markers, the values of empty ones were estimated as described in Supporting Information) for each consecutive step of filtration. **b** A control measurement of filtering the free dye (Alexa Fluor 488) in the absence of nanoparticles, performed in the same manner as in (a). **c** Normalized FCS curves (g(τ)) of TiO_2_ nanotubes, labelled with Alexa Fluor 488, before filtration (red), after filtration (green), and with the same filtered sample measured after 14 days (blue). Note that the FCS curve of nanomaterial before filtration (red) is almost undistinguishable from the curve of free dye (orange). **d** Comparison of confocal fluorescence micrographs of LA-4 cells (cell membranes labelled with CellMask Orange, shown in green) exposed to labelled TiO_2_ nanotubes with various amounts of free dye in the sample (the bound and unbound dye is Alexa Fluor 647, shown in red and scaled so its fluorescence intensity is directly comparable across the images): a combination of labelled nanomaterial and a 100-fold excess of free dye (300 nM), mimicking inefficient dye removal (left), cells exposed to correctly labelled and filtered nanomaterial (approximately 3 nM of bound dye, middle), and cells exposed to free dye (1.6 nM, left). The ratio between the surface of the nanomaterial and surface of cells is 0:1, 1:1 and 1:1 (from left to right). The localization of dye on the left two micrographs is very similar (arrowheads). For micrographs of separate color channels refer to Figure S20 in Supporting Information.

From the fluorescence signal of the final, cleaned retentate, we can estimate the degree of labelling of the nanomaterial. If the calculated number of dye molecules per nanoparticle settles at a value between 1 and 10, the nanoparticles are considered well labelled. In our case, the signal of the retentates settled at a value equivalent to the dye concentration of around 300 nM (orange symbols in Figure 3a), corresponding to 1.5 dye molecules per nanotube (calculations are discussed in Supporting Information). However, if the fluorescence of retentates does not settle (as demonstrated with filtration of a free dye in Figure 3b), the dye did not bind to the nanomaterial, indicating that a change in the labelling strategy is required.

In an ideal case with no desorption of the dye from the nanoparticles, the filtrates’ signal should decrease exponentially with each subsequent filtration (Figure 3b). In a more realistic scenario (Figure 3a), however, the amount of the dye in the filtrate stabilizes after a few filtrations due to steady desorption of the dye from the nanoparticles, with a lower value corresponding to more stable binding. Also, if the filtered sample is filtered again after a few days or weeks, the intensity of the first next filtrate will be higher than of the previous filtrate as it will contain all the dye that desorbed during this time period (Figure 3a, filtrates #6 and #7). Note that measuring retentates at each step of filtration includes pipetting of small quantities of inhomogeneous nanomaterial suspensions, which can decrease the precision of determined amount of dye in the retentate (hence the difference between retentates #6 and #6b in Figure 3a).

##### Evaluation by FCS

The nanoparticle labelling and free dye removal are additionally evaluated with fluorescence correlation spectroscopy (FCS) (Figure 3c), ^[56]^ which can distinguish free dye from the dye on the nanoparticles based on their diffusion properties, associated with their hydrodynamic radius. This can be determined via autocorrelated fluctuations of detected light, arising as the particles diffuse in and out of the observation volume ^[57,58]^ (similarly as in dynamic light scattering (DLS), but relying on fluorescently emitted rather than scattered photons). As demonstrated in Figure 3c, faster-diffusing, smaller entities with shorter transit times generate autocorrelation curves shifted towards shorter times (dotted curve, e.g. free dye) compared to larger, slower entities (dashed curve, e.g. dye on nanoparticles). From additional analysis of the autocorrelation curves and fluorescence fluctuations, one can also obtain the diffusion coefficient, size of particles, their concentration, fraction of unbound dye, and number of dyes per nanoparticle. ^[56]^

Note that FCS should be measured on retentates at a pH where the nanomaterial is stably dispersed. Otherwise, the light emitted from aggregates overwhelms the readouts of smaller diffusing molecules and nanoparticles, preventing the determination of diffusion properties of the latter. Importantly, such FCS measurements can be quickly performed even with a laser-scanning confocal fluorescence microscope just before an experiment to double-check the nanomaterial’s state. ^[59]^ As shown in Figure 3c, the FCS curves of the non-filtered sample (red, **iii**) are dominated by the signal of the free dye (orange). In contrast, a well-filtered sample contains much slower diffusing entities (green, **iv**), indicating that most of the dye diffuses together with nanoparticles. FCS curves of intermediate filtering retentates, as well as of an old sample with some dye desorbed, show a weighted sum of these two extreme cases (blue, **iv\***), indicating the presence of both free and bound dye (see also Figure S21 in Supporting Information).

#### 2.3.5. Potential Artefacts

To resume, unbound dye in the sample can arise from the following:

- insufficiently removed unbound excess dye from the labelling step,
- desorbed dye that was adsorbed on the nanoparticle surface during the labelling step,
- desorbed linker with covalently bound dye.

All of these processes contribute to the false colocalisation and misinterpretation of the measurements, when it is impossible to discern fluorescently labelled nanoparticles (Figure 3d, middle) from large quantities of free dye (Figure 3d, left)..

Based on the fluorescence intensity and FCS checkpoint, the linker can be optimally selected to minimize desorption from the nanomaterial. This procedure has enabled us to identify AEAPMS linker as the most suitable for reliable tracking of TiO_2_ nanotubes. Its desorption was found to be relatively slow, and well-filtered samples were stable in the buffer for a few days (Figure 3a). If the nanomaterial has been stored for extended periods, we filter the nanomaterial at least once again before any cell-exposure experiments. Note that even if some of the dye desorbs from otherwise well-cleaned nanomaterial, its signal is homogeneously distributed over the large sample volume and therefore negligible (Figure 3d, right) compared to the signal of a similar amount of the dye bound to the nanomaterial (Figure 3d, middle). Also, in our case only a small fraction of the Alexa Fluor 647 dye was found internalized and localized in vesicles in the cells after a 2-day incubation (Figure S18 Supporting Information).

### 2.4. Characterization of Labelled Nanomaterial

Once the labelling is successfully performed, it must be verified that the labelled material preserved the desired properties of the native material, such as its native charge and morphology, which both knowingly affect interactions with cells. ^[48,49,60–62]^ Otherwise the functionalization and labelling procedure must be appropriately adjusted.

#### 2.4.1. Potential Artefacts due to Altered Charge

The final surface charge of the labelled nanomaterial is influenced by the amounts of bound linker and dye. Labelling the same functionalized nanomaterial at different conditions (dye concentration, timing, buffer etc.) can result in different pH-dependencies of Zeta potential (Figure S9 in Supporting Information), which can lead to different cellular response, as demonstrated before (Figure 2d). For cell exposure experiments, we therefore selected the labelled nanomaterial whose Zeta potential measurements were relatively close to those of the native nanomaterial (Figure 2c, **iv**).

These measurements can also guide optimization of the protocol: in our case, both the native and labelled TiO_2_ nanotubes are stable in dispersion at pH 10 according to the measured Zeta potentials, so they are always sonicated and stored in 100-times diluted bicarbonate buffer with pH 10. The buffer is diluted to minimize screening of the charge on the nanomaterial by ions in the buffer, thus enhancing the electrostatic repulsion and preventing aggregation of nanoparticles.

#### 2.4.2. Potential Artefacts due to Altered Morphology

We next verified that the morphology of our TiO_2_ nanotubes, determined from TEM measurements, remained similar to that of the native nanomaterial (**Figure 4a**, top three rows) despite the 45-minute sonication used in the labelling protocol, most of which is employed to increase labelling efficiency. However, longer sonication times markedly changed the morphology of the nanotubes (Figure 4a, bottom). The TiO_2_ nanotubes that were shorter due to a 2 hour sonication (Figure 4b, bottom) also affected the cell’s morphology differently than nanotubes sonicated for 45 minutes (Figure 4b, middle), which showed similar cellular response as non-labelled nanotubes sonicated for only 15 min (Figure 4b, top). It is therefore essential to monitor for possible morphology changes of the nanomaterial, as they influence interactions with the cells and therefore the outcomes of cell-exposure experiments.

**Figure 4.**
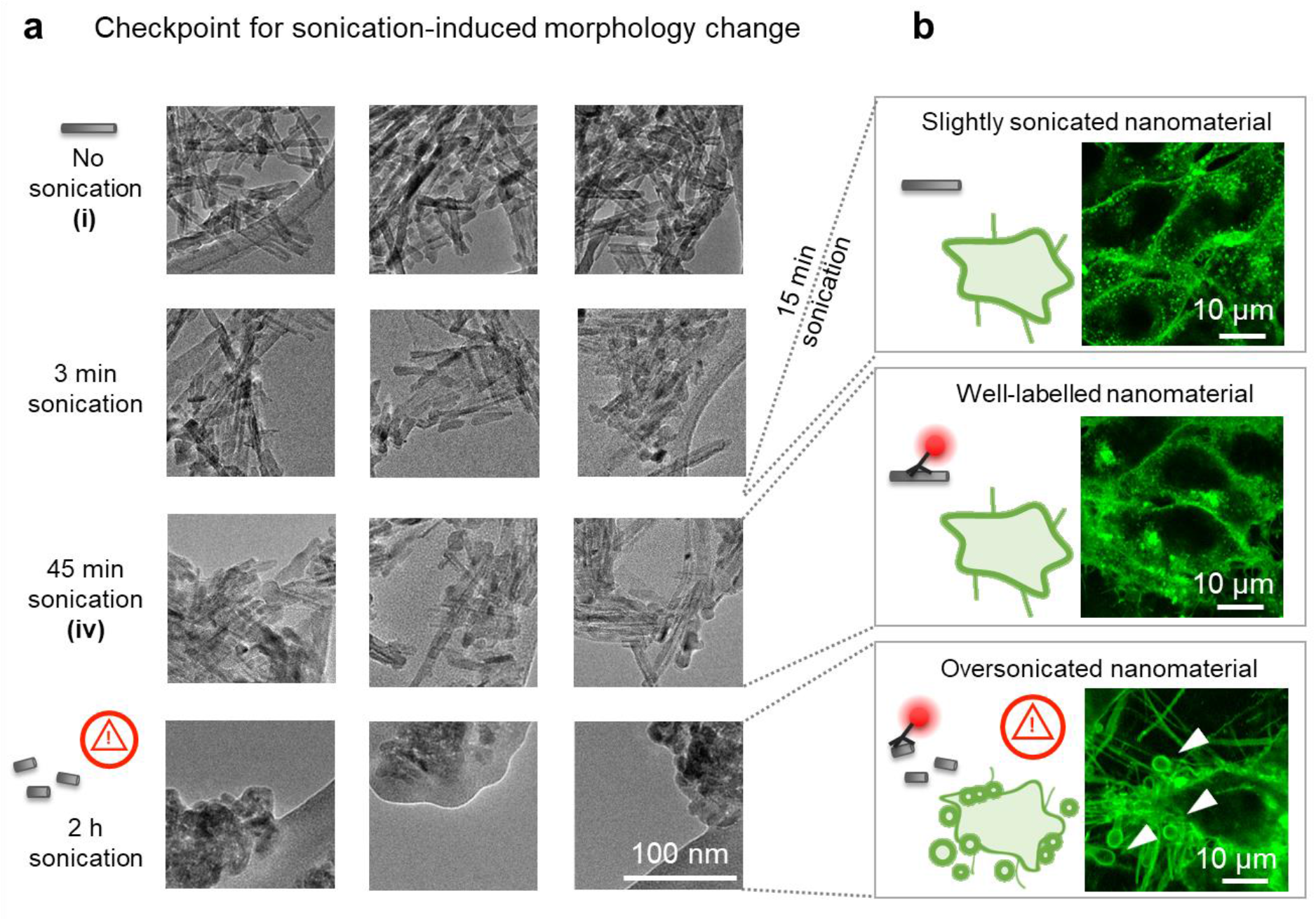
Inspection of the morphology of the nanomaterial to avoid sonication-induced modifications. **a** TEM images of (top to bottom) the initial nanoparticle sample, and nanoparticles sonicated for 3 min, 45 min and 2 h, with three sites per sample shown in columns. The delivered dose is 2 kJ per minute of sonication. The large flat structure in TEM images at 2 h sonication is a lacey carbon TEM grid. **b** Confocal fluorescent micrographs of cells (cell membranes are labelled with CellMask Orange and shown in green), exposed to TiO_2_ nanotubes sonicated for 15 minutes (top), 45 minutes (middle) and 2 h (bottom) with the ratio between the surface of the nanomaterial and surface of cells being 10:1, 1:1 and 1:1, respectively., The nanotubes in the lower two images are labelled with Alexa Fluor 647, the fluorescence of which is not shown. Note the vesicles and long filaments on the lowest micrographs (arrowheads) which are not observed in the other two micrographs.

#### 2.4.3. Other Potential Artefacts due to Sonication

It is also worth noting that many dispersion protocols use albumin or other proteins to improve the dispersion stability of nanomaterials. However, sonication may destabilize proteins and enhance their aggregation, which leads to the formation of biologically active, potentially immunogenic amyloid structures. ^[63]^ To exclude such effects, one could include a sham control to the experiments by sonicating the sonication medium without the nanoparticles and investigate its effects on the cells. If possible, however, a protocol for dispersing the nanomaterial without sonication of proteins is preferred.

Another often overlooked issue, related to tip-sonication of nanomaterials, is erosion of the transducer (tip) during sonication, which can contaminate the sample with metallic particles, ^[64]^ see also Figure S22 in Supporting Information. Tip erosion is increasingly prominent for longer sonication times, but can be mitigated by carefully polishing the tip and thoroughly washing it to remove debris from the polishing procedure. Tip polishing is best done between each of the subsequent sonications, or at the latest when a dark ring forms on the tip (Figure S22 in Supporting Information).

### 2.5. An Example of Detection of Early Molecular Events

After carefully following the labelling procedure with appropriate quality control steps, the well-characterized nanomaterial is ready for application in experiments. The rigorous evaluation of the material at all key preparation steps ensures minimal misinterpretation of the outcomes.

We exemplify the application of the resulting material in **Figure 5**, where cells have been incubated with TiO_2_ nanoparticles, labelled with Alexa Fluor 647, for two days before membrane staining and imaging. By using high-resolution fluorescence microscopy (stimulated emission depletion microscopy - STED), nanoparticle-cell interactions can be resolved with a resolution well below the diffraction limit (Figure 5c). Co-localization of both fluorescence signals indicates that the nanoparticles are wrapped with the cellular membrane, as we reported previously ^[17]^. In our further studies, advanced fluorescence microscopy with well-labelled nanomaterial enabled us to discover several other related events of interaction between nanomaterial and diverse cellular structures (**Figure 6**). Visualization and spatio-temporal characterization of such early molecular events allowed us to unravel causal relationships between them and devise their link to adverse outcomes after exposure to nanomaterials. In particular, we elucidated the mechanism how certain types of nanomaterials trigger previously unexplained chronic inflammation, based on which we designed an all-in-vitro test for its prediction. ^[19]^

**Figure 5:**
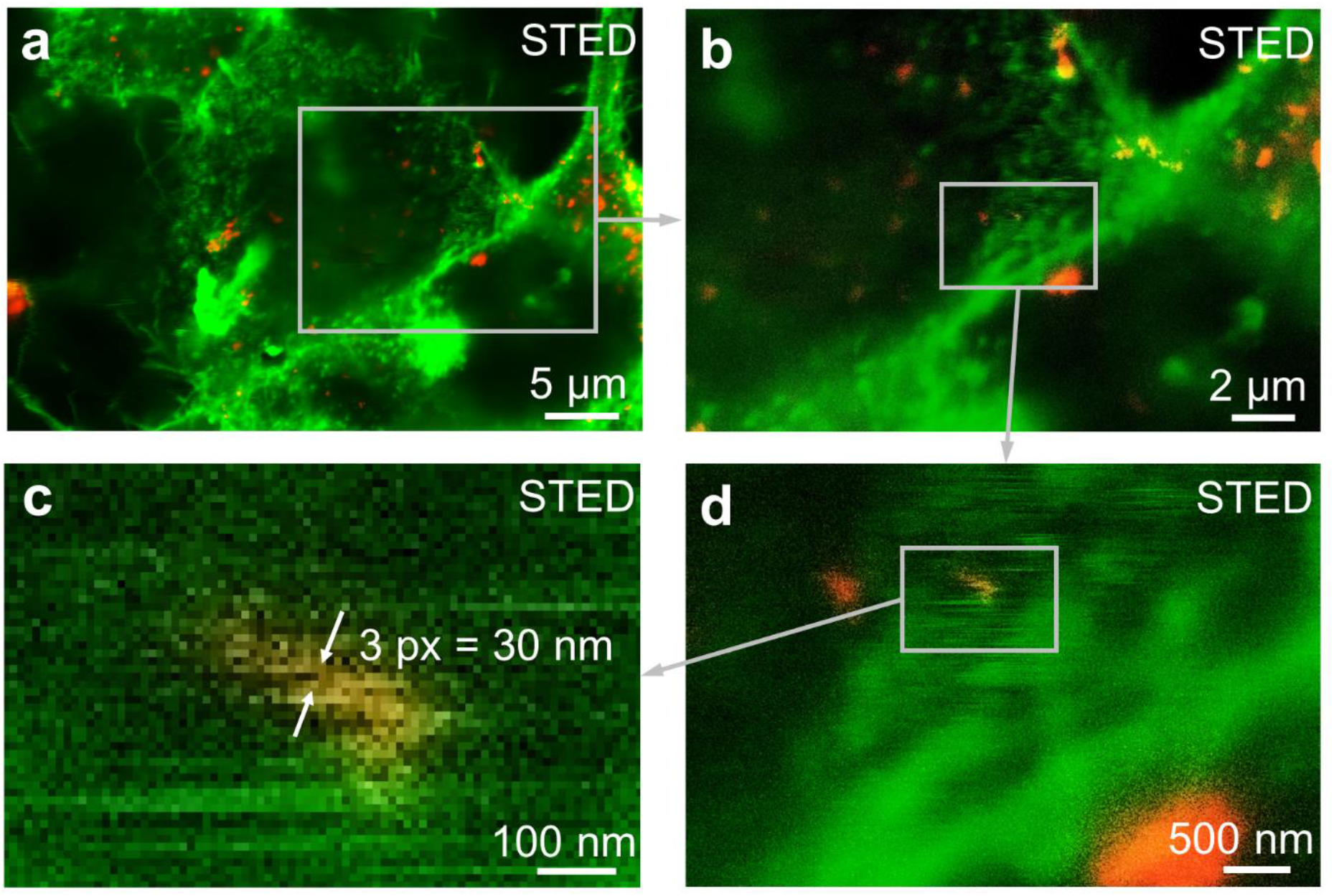
Super-resolution imaging of fluorescently labelled TiO_2_ nanotubes in living cells. **a** The original STED image, and **b–d** corresponding zoom-ins, of LA-4 cells (membranes labelled with CellMask Orange, green) incubated for 2 days with efficiently and stably labelled nanoparticles (Alexa Fluor 647, red), the ratio between the surface of the nanomaterial and surface of cells being 1:1, as an example of an in vitro experiment. The signal of nanomaterial is logarithmically scaled. For micrographs of separate color channels refer to Figure S23 in Supporting Information.

**Figure 6:**
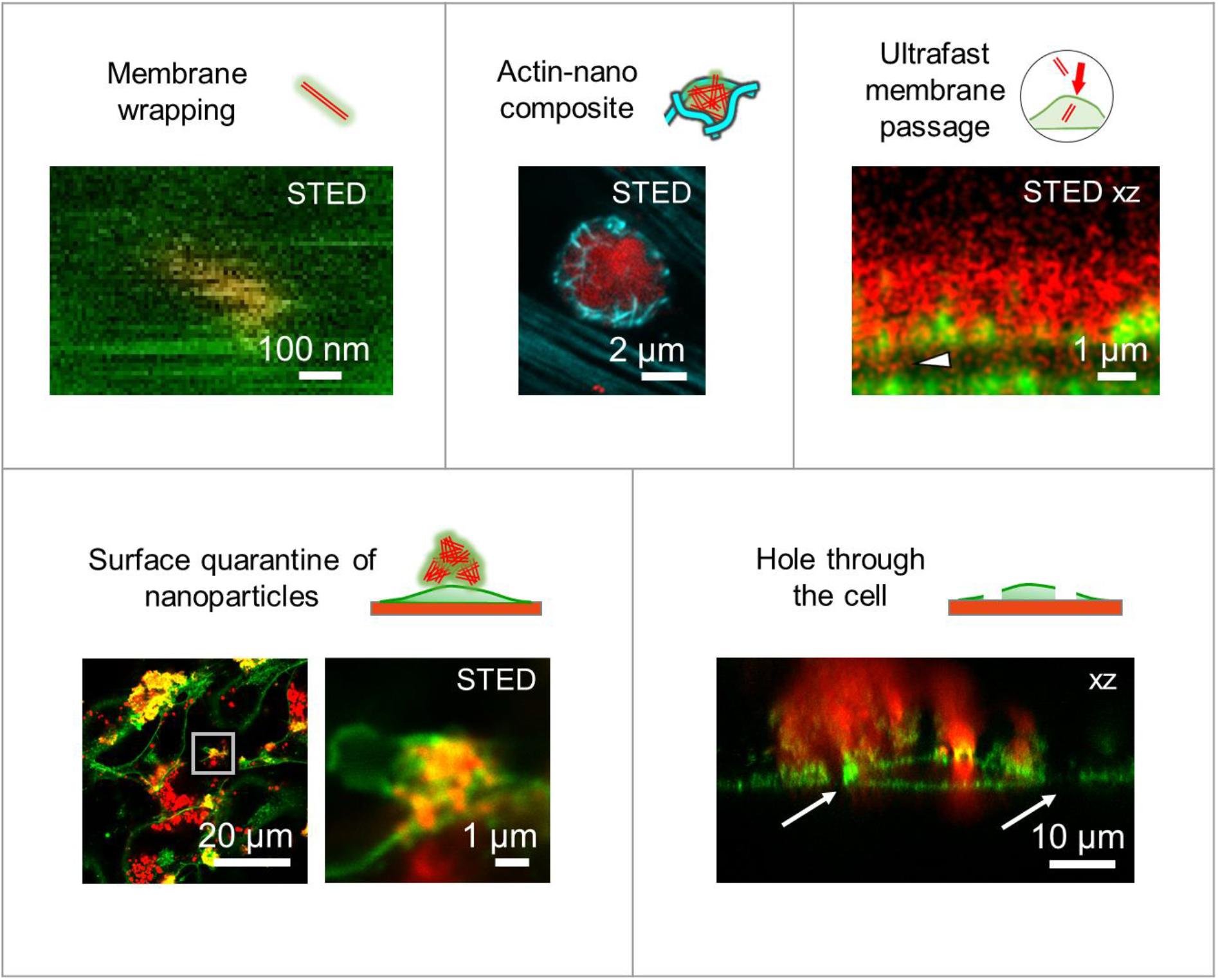
Examples of five early molecular events in living LA-4 cells following exposure to nanomaterial (TiO_2_ nanotubes labelled with Alexa Fluor 647, red), detected with confocal or STED fluorescence microscopy. Cell membranes are labelled with CellMask Orange (green), actin with SiR Actin (cyan). For micrographs of separate color channels refer to Figure S24 in Supporting Information.

## 3. Conclusion

The use of fluorescently labelled nanomaterial in live-cell fluorescence microscopy requires stable and reliable binding of fluorescent dyes to the nanomaterial of interest, which, noteworthy, can modify nanomaterial properties and influence experimental results. Thus, to ensure the relevance and reproducibility of the experimental studies, it is of utmost importance to thoroughly characterize the labelled nanomaterial’s state and properties.

We here presented a procedure for quality control of nanoparticle labelling, which utilizes the following steps:

- Comparison of nanomaterial morphology before and after sonication by TEM: prolonged sonication can result in altered size/shape of the nanomaterial, causing it to interact differently with the cells than the native nanomaterial.
- Determination of the nanomaterial’s surface charge before and after labelling by measuring the pH-dependency of Zeta potential: both the linker and dye influence the surface charge of the nanomaterial, affecting the response of cells as well.
- Removal of unbound dye by centrifugal filtration, monitored by intensity and FCS measurements: when free or desorbed dye is not sufficiently removed from the sample, the fluorescence signal can be falsely assigned to nanoparticles.

We illustrated the potential artefacts in cell-exposure experiments arising from missing out each step, and provided further practical advice on mitigating any issues should they arise. We here demonstrated the procedure to control the labelling of TiO_2_ nanotubes functionalized with silane linkers and labelled with SDP-/NHS-esters of Alexa Fluor dyes. The approach can easily be modified to ensure reliable labelling of other types of nanoparticles ^[45–47]^ and colloidal particles of up to 10 μm in size, above which the Zeta potential and fluorescence correlation spectroscopy experiments become unreliable.

By carefully monitoring the entire labelling procedure, identifying and troubleshooting unsuccessful labelling is significantly simplified. Moreover, the suggested procedure results in well-characterized and stably labelled nanomaterial that reproduces the pristine material’s mode-of-action, thus enabling reproducible, relevant and reliable nanotoxicity studies with advanced microscopies in live cells. These can uniquely identify and connect key molecular events that trigger adverse outcomes, which can lead towards better mechanistic understanding of nanoparticles’ effects on our health, design of more efficient testing strategies, and finally, safer use of the impressive technologies the nanomaterials can offer.

## 4. Experimental Methods

### Materials

Alexa Fluor 488 SDP ester (Thermo Fisher), Alexa Fluor 647 NHS ester (Thermo Fisher), CellMask Orange (Invitrogen), LCIS-Live Cell Imaging Solution (Invitrogen), #1.5H µ-dishes (Ibidi), PBS – phosphate buffer saline (Gibco), 100x dcb: 100-times diluted bicarbonate buffer (pH 10, osmolarity 5 mOsm L^-1^, mixed in-house), F-12K cell culture medium (Gibco), Trypsin (Sigma), FBS (Fetal bovine serum, ATCC), Penicillin-Streptomycin (Sigma), Non-essential amino acids (Gibco), BSA-bovine serum albumin (Sigma), 3-(2-aminoethylamino)propyltrimethoxysilane (AEAPMS, Alfa Aesar), lacy carbon film supported by a 300-mesh copper grid (Ted Pella, Inc.), sodium bicarbonate buffer mixed in-house from NaHCO3 (Merck) and Na2CO3 (Kemika), anhydrous DMSO (Sigma Aldrich), ethanol (Carlo Erba), centrifugal filter device Amicon Ultra 4 mL 100K (Merck Millipore), HCl (Merck), KOH (Sigma Aldrich), KCL (Merck), polystyrene cuvettes for Zeta potential measurements (Sarstedt), 96-well black flat-bottom microplate (Brand)

### Software

Imspector (version 16.2.8282-metadata-win64-BASE) software provided by Abberior, SymPhoTime64 (PicoQuant), Mathematica 12.0 - license L5063-5112 (Wolfram)

### TiO_2_ Nanotubes Synthesis

TiO_2_ nanotubes were synthesized in-house using the method, described elsewhere. ^[22,65]^ Synthesis of the anatase TiO_2_ nanotubes with a diameter of 10 nm, mean length of 200 nm and a BET surface of 150 m^2^ g^−1^ proceeds in several stages:

1) Synthesis of sodium titanate nanotubes (NaTiNTs);

TiO_2_ + NaOH(aq) → (Na,H)_2_Ti_3_O_7_ xH_2_O

NaTiNTs are synthesized according to the method already reported elsewhere. ^[66]^ In short, a suspension of TiO_2_ (Titanium(IV) dioxide, anatase, 325 mesh, Aldrich) and 10 M NaOH (1 g TiO_2_ per 10 mL 10 M NaOH) is ultrasonicated for 30 min and then stirred for 1 hour at room temperature. The suspension is transferred to a Teflon-lined autoclave; the filling volume is 80%. The closed autoclave is heated for three days at 135 °C. When the reaction mixture is cooled down to room temperature, the resulting material is washed twice with deionized water (2 x 300 mL), once with ethanol, and dried overnight at 100 °C. At this stage morphology is checked with scanning electron microscope (SEM).

2) Transformation of sodium titanate nanotubes to hydrogen titanate nanotubes (H_2_Ti_3_O_7_); (Na,H)_2_Ti_3_O_7_ → H_2_Ti_3_O_7_ xH_2_O

Next, sodium ions are removed by an ion-exchange process; 2.5 g of sodium titanate nanotubes is dispersed into 300 mL of 0.1 M HCl(aq). ^[67]^ The prepared dispersion was stirred at room temperature for 1 h, centrifuged to separate the solid material from the solution. The dispersing, centrifuging and separating steps were repeated for three more times. At the end, the solid material is washed three times with 200 mL of distilled water and once with 100 mL of ethanol and dried overnight at 100 °C. At this stage the content of sodium and chlorine is determined with energy-dispersive X-ray spectroscopy (EDXS). If the content of sodium and/or chlorine is above 0.3 wt. % is the washing procedure with 0.1 M HCl and deionized water was repeated.

3) Transformation of hydrogen titanate nanotubes to TiO_2_ nanotubes; H_2_Ti_3_O_7_ xH_2_O → TiO_2_ + H_2_O

Finally, hydrogen titanate nanotubes are transformed to TiO_2_ nanotubes by thermal treatment in air. In general, 500 mg of H_2_Ti_3_O_7_ nanotubes is weighed in an alumina boat, placed into an oven and heated at a ramp rate of 1 °C min^-1^ to 380 °C (or 400 °C). The samples kept at 380 °C for 6 h, and cooled down to room temperature afterwards. ^[22]^ At this stage morphology is checked with SEM and TEM, structural analysis with XRD (X-ray powder diffraction), elemental composition and specific surface are determined.

### TiO_2_ Nanotubes Functionalization (f-TiO_2_)

TiO_2_ nanotubes (100 mg) are dispersed sonically in 30 mL of dry toluen and then heated to 60 °C while constantly stirring. Separately, a solution of 840 μL of 3-(2-aminoethylamino)propyltrimethoxysilane (AEAPMS, Figure S2 in Supporting information) in 30 mL of dry toluene is prepared. The solution of AEAPMS in toluene is added dropwise (30 minutes) to the dispersion of TiO_2_ nanotubes in toluene. After 16 h the reaction mixture is cooled to room temperature and the solid material is separated from the liquid phase by centrifugation. Then, it is rinsed first with 30 mL of toluene (3 times) and then 3 times with 30 mL of hexane. The isolated product is dried overnight at 80 °C and for 2 hours at a vacuum drier at 80 °C and 150 mbars. The quality of functionalization is checked with FTIR and Zeta potential measurements. A protocol for the functionalization is described in the Supporting Information.

### TiO_2_ Nanotubes Labelling

All labelling quantities refer to labelling of 1 mg of TiO_2_ nanotubes, functionalized with AEAPMS (f-TiO_2_), a linker with a positive charge at labelling pH. First, 1 mg of functionalized TiO_2_ (Excellence Plus, Mettler Toledo) is weighed and diluted in 1 mL of 100-times diluted sodium bicarbonate buffer with pH = 10.00, which has been prepared in a glass flat bottom flask with fresh miliQ water. The sample is transferred to a glass vial. Nanoparticles are then sonicated with a tip sonicator (MISONIX Ultrasound Liquid Processor with 419 MicrotipTM), with the average power 35 W (amplitude set to 70) and a microtip with radius of 3 mm with a surface of 30 mm^2^ for t_run_ = 15 min, t_on_ = 5 s, t_off_ = 5 s. This totals the sonication time to t = 15 min and total sonication dose 31.7 kJ.

After the first sonication, 3.2 μL of 12.1 mM fresh Alexa Fluor 488 or 647 – SDP ester (Life Technologies, Alexa Fluor 488 or Alexa Fluor 647, Figure S2 in Supporting information), diluted in anhydrous DMSO, is added. Due to linker being positively charged at labelling pH, a negatively charged dye is chosen. For each mg of nanomaterial 40 nmol of label is added. Again, the sonication is performed with a tip sonicator for t_run_ = 30 min, t_on_ = 5s, t_off_ = 5s with total sonication dose 31.7 kJ. This totals the cumulative sonication time to t = 45 min and total sonication dose of 95 kJ. Note that the sonication dose at which the nanomaterial morphology remains unaltered needs to be verified for each nanomaterial separately.

After the sonication is complete, a mixing magnet is added to the glass vial containing the sample and is left to stir overnight on a magnetic mixer at 260 rpm, covered with aluminum foil to prevent photo-bleaching of the dye. After approximately 12 – 16 h of incubation, the magnet is removed from the glass vial. The sample is then centrifuged to remove excess unbound label. A detailed protocol for the labelling is described in the Supporting Information.

### Free Dye Removal

The free dye is removed by centrifugation in a centrifugal filter device (CFD), which forces the sample through a filter during centrifugation, separating the sample into the filtrate below the filter (containing unbound dye) and retentate on the filter (containing nanoparticles, which are too large to pass through the filter), which is then resuspended. To ensure efficient filtration, the size of pores on the filter of the centrifugal device should be significantly larger than the fluorescent dyes and smaller than the nanoparticles.

Centrifugation is performed in a centrifuge filter device (CFD) (Amicon Ultra 4 mL 100K) with a 100 kDa membrane on centrifuge (LC-320, Tehtnica, Zelezniki) at 3000 rpms to achieve 1000 g. Firstly, the CFD is prepared for use according to the manufacturer’s guidelines, i.e. by centrifuging 2 mL of deionized water through the membrane. Then, 1 mL of labelled TiO_2_ nanoparticles are pipetted into the CFD and spun for 15 minutes. Afterwards, the filtrate is stored for further analysis and the retentate is resuspended (by mixing with a pipette) in 1 mL of medium, consisting of 70% ethanol and 100-times diluted bicarbonate buffer (39.37 mL of 96% ethanol is mixed with 0.5 mL non-diluted bicarbonate buffer and 11.27 mL of deionized water). The empty filtrate compartment is filled with the same medium so the membrane is submerged (this is to ensure good contact in the ultrasonic bath in the next step). The CFD is sonicated on the ultrasound bath (Branson 2510) for 20 seconds to diminish the amount of nanoparticles stuck to the CFD membrane and to break up the nanoparticle aggregates. The filtrate compartment is washed with ethanol and dried to prevent mixing of subsequent filtrates, and the sample in the CFD is spun again.

The sample is filtered twice in this manner and two times more with 100-times diluted bicarbonate buffer as medium instead of the mixture of 70% ethanol and 100-times diluted bicarbonate buffer to reduce the amount of ethanol in the sample. As a general guideline, the filtered sample should have a noticeable color at 1 mg/mL, as should the very first filtrate and the color should decrease in subsequent filtrates (Figure S11 and Figure S12, Supporting information). It is also worth noting that a general idea of the cleaning efficiency can already be obtained by measuring the original sample, final cleaned sample and last filtrate (or, even better, two last filtrates). The sample should be filtered at least once more just prior to the experiments in the desired media, if it was stored in a refrigerator for longer periods of time (in our case more than two weeks). A detailed protocol for the filtration process is described in the Supporting Information.

### TEM and FTIR

The morphology of nanomaterials was investigated with a transmission electron microscope (TEM Jeol 2100, 200 keV). Specimens of nanoparticles were dispersed ultrasonically in methanol and a drop of the dispersion was deposited onto a lacy carbon film supported by a 300-mesh copper grid. Infrared spectra were obtained via Attenuated total reflection Fourier transform infrared spectroscopy (ATR-FTIR) with a universal ATR accessory on a Spectrum 400 FTIR spectrometer (Perkin Elmer). The spectra were recorded with the resolution of 4 cm^-1^ and 16 scans.

### Zeta Potential Measurements

Zeta potential measurements were performed on NanoBrook ZetaPALS Potential Analyzer (Brookhaven) and electrode for non-organic solvents (AQ-1203, Brookhaven). For measurement of each part of the Zeta potential, acid and base, 3 mL of sample diluted to c = 0.1 mg mL^-1^ in 10 mM KCl was used. The pH was measured with a pH meter (Seven Multi, Mettler Toledo) and 1 mm thick pH electrode (Inlab ExpertPro, Mettler Toledo) right before every measurement of the Zeta potential. Each one of 10 measurements at each pH value was measured until the experimental fit errors (residuals) were below the set value of 0.025. From the 10 measurements, the average and error were calculated.

The change of pH was achieved by addition of HCl (to lower pH) or KOH (to rise pH). After each addition, the sample was left on a magnetic stirrer (IKA topolino) for approximately 3-5 min or until stabilization of pH occurred.

### Fluorescence Intensity of Filtrates

Free dye concentration measurements were performed on the spectrometer Tecan Infinite M1000 (Tecan Group Ltd., Männedorf, Switzerland). Samples have been loaded to 96-well plates (96-well plate, Brand) prior measurements in volumes of 150 μL. The amount of dye in the sample was calculated from the known volumes and concentrations, which were calculated from the measured fluorescent signal and previously measured calibration curves for concentration, medium and nanomaterial concentration in the sample. Direct comparison between samples was possible since all experimental conditions were the same (volume of sample, temperature, machine settings, etc.).

The filtrates were measured as-is, whereas a fraction of each retentate was taken away after the filtration and diluted to achieve the 150 μL needed for the fluorescence measurement in order to not use too much of the valuable labelled nanomaterial. For the same reason, fluorescence of intermediate retentates was estimated from the measured intensities of filtrates and other retentates (open circles in Figure 3a). Due to variations in filtration volumes, we calculated the absolute quantities of dye in the whole sample including the retentates used for measurements. A more detailed description is located in the Supporting Information.

### Cell Culture and Sample Preparation for Microscopy

All fluorescence microscopy was done on LA-4 murine lung epithelial cells which were cultured according to ATCC guidelines: they were grown in a mixture of F-12K medium (Gibco), 15% FBS (Fetal bovine serum, ATCC), 1% P/S (Penicillin-Streptomycin, Sigma), 1% NEAA (non-essential amino acids, Gibco) in an incubator at 37 °C, saturated humidity and 5% CO_2_. After incubation with nanomaterial, the culture conditions remained the same.

For the fluorescence image comparison in Figure 1-4, the LA-4 cells were seeded in a 1.5H Ibidi µ-dish at 30% confluency. Two days later, the appropriate sample of nanomaterial was added to the cells. For “well-labelled nanomaterial” (Figure 2d left, Figure 3d middle, and Figure 4b above), 35 µL of 0.1mg mL^-1^ TiO_2_ nanotubes fluorescently labelled with Alexa Fluor 647 in PBS (phosphate-buffered saline) were added to 315 µL of full cell medium on cells to achieve a 1:1 ratio between the surface of the nanomaterial and surface of cells (so-called surface dose). For sample preparation of other, incorrectly labelled nanomaterial, see below. After 2 days of incubation with the nanomaterial, the medium with cells was removed and cells were labelled with 100 µL of 5 µg mL^-1^ CellMask Orange in Live Cell Imaging Solution (LCIS, Molecular Probes) for 6 minutes at 37 °C. Later, the medium with CellMask Orange was slowly removed and cells were observed in 100 µL LCIS.

The “native nanomaterial” in Figure 2d (left) was incubated with a 10:1 surface dose of nonlabelled nanomaterial, the “nanomaterial with different surface charge” in Figure 2d (middle) was incubated with a 1:1 surface dose of functionalized TiO_2_ nanotubes. The “desorbed fluorophores” in Figure 3d (left) was incubated with a 1.6 nM concentration of free Alexa Fluor 647, the “unwashed label” in Figure 3d (right) was incubated with a 1:1 surface dose of well-labelled TiO_2_-Alexa647 and free Alexa Fluor 647 to achieve a 3 nM final concentration of Alexa Fluor 647 on the nanomaterial and an additional 300 nM concentration of unbound Alexa Fluor 647, mimicking insufficient filtration. The “slightly sonicated nanomaterial” shown in Figure 4b (above) was sonicated for 15 minutes and exposed to the cells at a 10:1 surface dose, and the “oversonicated” nanomaterial shown in Figure 4b (below) was sonicated for 2 hours instead of 45 minutes during the labelling procedure and exposed to the cells at a 1:1 surface dose.

For the high-resolution STED image, the LA-4 murine lung epithelial cells were seeded into a 1.5H Ibidi µ-Dish at 30% confluency and cultured in complete medium. A 100 µg mL^-1^ dispersion of TiO_2_ nanotubes labelled with Alexa Fluor 647 in PBS was diluted in 270 µL of full cell medium to achieve the final concentration of 10 µg mL^−1^. When added to the cells, this corresponded to a surface dose of 1:1. After 2 days of incubation, the samples were washed and the plasma membranes were labelled with 100 µl of 5 µg mL^−1^ CellMask Orange in PBS. The sample was washed with PBS after 20 minutes of incubation and observed in PBS right afterwards. Larger confocal fields of view can be found in Figure S25 in Supporting information.

### FCS

For Fluorescence Correlation Spectroscopy (FCS), a PicoQuant MicroTime 200 confocal microscope was used. An Olympus IX81 inverted microscope was equipped with a 60x water immersion objective (Olympus UPLSAPO 60xW NA1.2). Alexa Fluor 488 was excited by a pulsed laser of wavelength 488 nm (PicoQuant, Germany), at 40 MHz repetition rate and average power of about 5 μW at the objective. Fluorescence of the dye was acquired through a 531/40 emission filter (Brightline, Semrock) by an APD detector (tau-SPAD, PicoQuant, Germany), connected to a PicoQuant HydraHarp 400 single-photon-counting unit. Typical acquisition times were 5–15 min. The fluorescence fluctuations were analyzed using SymPhoTime64 (PicoQuant). For presentation, FCS curves were normalized to ease the comparison of their relative shapes.

### Fluorescence Microscopy and STED

The imaging was performed with an Abberior Instruments STED microscope using a 60x water immersion objective (Olympus UPLSAPO 60xW NA1.2) and the accompanying software Imspector (version 16.3.11462-metadata-win64). The fluorescence of CellMask Orange and TiO_2_-Alexa647 was excited using two pulsed lasers at 561 and 640 nm, and their fluorescence was detected by two avalanche photodiodes at 580– 625 nm and 655–720 nm (filters by Semrock), respectively.

The confocal fluorescence images in Figure 1–4 were acquired using a pixel size of 100 nm, dwell-time 10 µs, and laser powers at around 10 µW.

The STED image in Figure 5 was acquired using a 10-nm pixel size. The resolution was increased by stimulated depletion of fluorescence by an 80 mW doughnut-shaped STED laser beam at 775 nm.

All experiments were performed at room temperature.

## Supporting Information

Supporting Information can be accessed in the associated pdf file.

## Acknowledgements

This research was funded by EU Horizon2020 Grant No. 686098 (SmartNanoTox project), Slovenian Research Agency (program P1-0060), Young Researcher Program (PR-08331, H. Kokot), Marie Skłodowska-Curie Individual Fellowship (grant no. 707348 to I. Urbančič), and Wolfson Foundation / Medical Research Council (MRC, grant no. MC_UU_12010/unit programmes G0902418 and MC_UU_12025) / Deutsche Forschungsgemeinschaft (Jena Excellence Cluster “Balance of the Microverse”, Collaborative Research Center 1278 “Polytarget”) to C. Eggeling.

The authors thank Monika Koren, Neža Golmajer Zima, Petra Čotar and David Dolhar for their help with the preparation of samples, Aleksandar Sebastijanović for kindly providing the micrograph of actin-nano composite as well as drs. Silvia Galiani and Falk Schneider for their help with the FCS experiments.

Boštjan Kokot and Hana Kokot contributed equally as first authors, and Tilen Koklič, Iztok Urbančič, and Janez Štrancar contributed equally as corresponding authors.

BK, HK, PU, SP, MG, CE, TK, IU, JŠ designed the study and analysis.

PU synthesized the nanoparticles.

PU and SP functionalized the nanoparticles.

BK, HK, KPvM labelled the nanoparticles.

BK, HK, PU, KPvM, IU prepared the samples.

BK, HK, PU, IU performed the experiments.

BK, HK, IU analyzed the data.

CE, TK, IU, JŠ supervised the study.

BK, HK prepared the manuscript with input from all other authors: PU, KPvM, SP, MG, CE, TK, IU, JŠ.

The authors declare no competing interests.

## References

[1] S. J. Klaine, P. J. J. Alvarez, G. E. Batley, T. F. Fernandes, R. D. Handy, D. Y. Lyon, S. Mahendra, M. J. McLaughlin, J. R. Lead, Environ. Toxicol. Chem. 2008, 27, 1825.

[2] P. Biswas, C.-Y. Wu, J. Air Waste Manag. Assoc. 2005, 55, 708.

[3] R. D. Handy, R. Owen, E. Valsami-Jones, Ecotoxicology 2008, 17, 315.

[4] S. Senapati, A. K. Mahanta, S. Kumar, P. Maiti, Signal Transduct. Target. Ther. 2018, 3, 1.

[5] D. Ziental, B. Czarczynska-Goslinska, D. T. Mlynarczyk, A. Glowacka-Sobotta, B. Stanisz, T. Goslinski, L. Sobotta, Nanomaterials 2020, 10, 387.

[6] J.-H. Lim, D. Bae, A. Fong, J. Agric. Food Chem. 2018, 66, 13533.

[7] Z. Chen, S. Han, S. Zhou, H. Feng, Y. Liu, G. Jia, NanoImpact 2020, 18, 100224.

[8] R. Gupta, H. Xie, J. Environ. Pathol. Toxicol. Oncol. Off. Organ Int. Soc. Environ. Toxicol. Cancer 2018, 37, 209.

[9] E. Baranowska-Wójcik, D. Szwajgier, P. Oleszczuk, A. Winiarska-Mieczan, Biol. Trace Elem. Res. 2020, 193, 118.

[10] A. Kroll, M. H. Pillukat, D. Hahn, J. Schnekenburger, Arch. Toxicol. 2012, 86, 1123.

[11] A. Nel, T. Xia, H. Meng, X. Wang, S. Lin, Z. Ji, H. Zhang, Acc. Chem. Res. 2013, 46, 607.

[12] C. J. Murphy, A. M. Vartanian, F. M. Geiger, R. J. Hamers, J. Pedersen, Q. Cui, C. L. Haynes, E. E. Carlson, R. Hernandez, R. D. Klaper, G. Orr, Z. Rosenzweig, ACS Cent. Sci. 2015, 1, 117.

[13] K. Gerloff, B. Landesmann, A. Worth, S. Munn, T. Palosaari, M. Whelan, Comput. Toxicol. 2017, 1, 3.

[14] S. Halappanavar, S. van den Brule, P. Nymark, L. Gaté, C. Seidel, S. Valentino, V. Zhernovkov, P. Høgh Danielsen, A. De Vizcaya, H. Wolff, T. Stöger, A. Boyadziev, S. S. Poulsen, J. B. Sørli, U. Vogel, Part. Fibre Toxicol. 2020, 17, 16.

[15] K. T. Thurn, T. Paunesku, A. Wu, E. M. B. Brown, B. Lai, S. Vogt, J. Maser, M. Aslam, V. Dravid, R. Bergan, G. E. Woloschak, Small 2009, 5, 1318.

[16] K. Kenesei, K. Murali, Á. Czéh, J. Piella, V. Puntes, E. Madarász, J. Nanobiotechnology 2016, 14, 55.

[17] I. Urbančič, M. Garvas, B. Kokot, H. Majaron, P. Umek, H. Cassidy, M. Škarabot, F. Schneider, S. Galiani, Z. Arsov, T. Koklic, D. Matallanas, M. Čeh, I. Muševič, C. Eggeling, J. Štrancar, Nano Lett. 2018, 18, 5294.

[18] S. Chen, J. Wang, B. Xin, Y. Yang, Y. Ma, Y. Zhou, L. Yuan, Z. Huang, Q. Yuan, Anal. Chem. 2019, 91, 5747.

[19] H. Kokot, B. Kokot, A. Sebastijanović, C. Voss, R. Podlipec, P. Zawilska, T. Berthing, C. B. López, P. H. Danielsen, C. Contini, M. Ivanov, A. Krišelj, P. Čotar, Q. Zhou, J. Ponti, V. Zhernovkov, M. Schneemilch, Z. M. Doumandji, M. Pušnik, P. Umek, S. Pajk, O. Joubert, O. Schmid, I. Urbančič, M. Irmler, J. Beckers, V. Lobaskin, S. Halappanavar, N. Quirke, A. Lyubartsev, U. Vogel, T. Koklič, T. Stoeger, J. Štrancar, Adv. Mater. 2020, DOI 10.1002/adma.202003913.

[20] T. Xia, M. Kovochich, M. Liong, L. Mädler, B. Gilbert, H. Shi, J. I. Yeh, J. I. Zink, A. E. Nel, ACS Nano 2008, 2, 2121.

[21] T. Baumgärtel, C. von Borczyskowski, H. Graaf, Beilstein J. Nanotechnol. 2013, 4, 218.

[22] M. Garvas, A. Testen, P. Umek, A. Gloter, T. Koklic, J. Strancar, PLOS ONE 2015, 10, e0129577.

[23] K. T. Ranjit, I. Willner, S. H. Bossmann, A. M. Braun, Environ. Sci. Technol. 2001, 35, 1544.

[24] M. Bettinelli, A. Speghini, D. Falcomer, M. Daldosso, V. Dallacasa, L. Romanò, J. Phys. Condens. Matter 2006, 18, S2149.

[25] P. Pietzonka, B. Rothen-Rutishauser, P. Langguth, H. Wunderli-Allenspach, E. Walter, H. P. Merkle, Pharm. Res. 2002, 19, 595.

[26] S. Grafmueller, M. P, D. L, M. LD. Pa, H.H, J. W, K. Hf, B.-T.T von M.U, W. P, Sci. Technol. Adv. Mater. Sci. Technol. Adv. Mater. 2015, 16, 16, 044602.

[27] Y. Sahoo, A. Goodarzi, M. T. Swihart, T. Y. Ohulchanskyy, N. Kaur, E. P. Furlani, P. N. Prasad, J. Phys. Chem. B 2005, 109, 3879.

[28] K. Brown, T. Thurn, L. Xin, W. Liu, R. Bazak, S. Chen, B. Lai, S. Vogt, C. Jacobsen, T. Paunesku, G. E. Woloschak, Nano Res. 2018, 11, 464.

[29] Y. Ge, Y. Zhang, S. He, F. Nie, G. Teng, N. Gu, Nanoscale Res. Lett. 2009, 4, 287.

[30] V. Holzapfel, A. Musyanovych, K. Landfester, M. R. Lorenz, V. Mailänder, Macromol. Chem. Phys. 2005, 206, 2440.

[31] X. Jiang, A. Musyanovych, C. Röcker, K. Landfester, V. Mailänder, G. U. Nienhaus, Nanoscale 2011, 3, 2028.

[32] L. Shang, L. Yang, F. Stockmar, R. Popescu, V. Trouillet, M. Bruns, D. Gerthsen, G. U. Nienhaus, Nanoscale 2012, 4, 4155.

[33] S. Giordani, J. Bartelmess, M. Frasconi, I. Biondi, S. Cheung, M. Grossi, D. Wu, L. Echegoyen, D.F. O’Shea, J. Mater. Chem. B 2014, 2, 7459.

[34] L. Shang, K. Nienhaus, X. Jiang, L. Yang, K. Landfester, V. Mailänder, T. Simmet, G. U. Nienhaus, Beilstein J. Nanotechnol. 2014, 5, 2388.

[35] S. Shahabi, L. Treccani, K. Rezwan, J. Nanoparticle Res. 2016, 18, 28.

[36] F. Hennrich, R. Krupke, K. Arnold, J. A. Rojas Stütz, S. Lebedkin, T. Koch, T. Schimmel, M. M. Kappes, J. Phys. Chem. B 2007, 111, 1932.

[37] C. M. Blumenfeld, B. F. Sadtler, G. E. Fernandez, L. Dara, C. Nguyen, F. Alonso-Valenteen, L. Medina-Kauwe, R. A. Moats, N. S. Lewis, R. H. Grubbs, H. B. Gray, K. Sorasaenee, J. Inorg. Biochem. 2014, 140, 39.

[38] W.-S. Cho, R. Duffin, F. Thielbeer, M. Bradley, I. L. Megson, W. MacNee, C. A. Poland, C. L. Tran, K. Donaldson, Toxicol. Sci. 2012, 126, 469.

[39] J. W. Soares, D. M. Steeves, D. Ziegler, B. S. DeCristofano, in Proc SPIE, 2006, pp. 637011-637011–9.

[40] J. Singh, J. Im, J. E. Whitten, J. W. Soares, A. M. Meehan, D. M. Steeves, in Proc SPIE (Eds.: Z. Gaburro, S. Cabrini, D. Talapin), 2008, p. 70300T.

[41] Q. Chen, N. L. Yakovlev, Appl. Surf. Sci. 2010, 257, 1395.

[42] J. Zhao, M. Milanova, M. M. C. G. Warmoeskerken, V. Dutschk, Colloids Surf. Physicochem. Eng. Asp. 2012, 413, 273.

[43] M. S. Killian, Organic Modification of TiO2 and Other Metal Oxides with SAMs and Proteins a Surface Analytical Investigation, PhD, 2013.

[44] T. Tenuta, M. P. Monopoli, J. Kim, A. Salvati, K. A. Dawson, P. Sandin, I. Lynch, PLOS ONE 2011, 6, e25556.

[45] J. R. LaGraff, J. C. Carnahan, D. Sangeeta, J. A. Ruud, Fluorescent Labeling Method and Substrate, 2003, US6607918 B2.

[46] J. A. Belmont, C. Corporation, Compositions Comprising Silane Modified Metal Oxides, 2010.

[47] S. Mallakpour, M. Madani, Prog. Org. Coat. 2015, 86, 194.

[48] A. Verma, F. Stellacci, Small 2010, 6, 12.

[49] M. Zhu, G. Nie, H. Meng, T. Xia, A. Nel, Y. Zhao, Acc. Chem. Res. 2013, 46, 622.

[50] D. Demandolx, J. Davoust, J. Microsc. 1997, 185, 21.

[51] T. Zimmermann, in Microsc. Tech., Springer, Berlin, Heidelberg, 2005, pp. 245–265.

[52] M. Gavrilovic, C. Wählby, J. Microsc. 2009, 234, 311.

[53] M. M. Frigault, J. Lacoste, J. L. Swift, C. M. Brown, J Cell Sci 2009, 122, 753.

[54] L. Zanetti-Domingues, T. Cj, R. Dj, C. Dt, M.-F. M, PloS One PLoS ONE 2013, 8, 8, e74200.

[55] L. Hughes, R. Rj, B. Sg, PLOS ONE 2014, 9, e87649.

[56] P. Rigler, W. Meier, J. Am. Chem. Soc. 2006, 128, 367.

[57] D. Magde, E. Elson, W. W. Webb, Phys. Rev. Lett. 1972, 29, 705.

[58] E. L. Elson, Biophys. J. 2011, 101, 2855.

[59] G. U. Nienhaus, P. Maffre, K. Nienhaus, in Methods Enzymol. (Ed.: S.Y. Tetin), Academic Press, 2013, pp. 115–137.

[60] O. R. Vasile, I. Serdaru, E. Andronescu, R. Truşcă, V. A. Surdu, O. Oprea, A. Ilie, B. Ş. Vasile, Comptes Rendus Chim. 2015, 18, 1335.

[61] S. Pujari-Palmer, X. Lu, M. K. Ott, Nanomaterials 2017, 7, DOI 10.3390/nano7040089.

[62] P. H. Danielsen, K. B. Knudsen, J. Štrancar, P. Umek, T. Koklič, M. Garvas, E. Vanhala, S. Savukoski, Y. Ding, A. M. Madsen, N. R. Jacobsen, I. K. Weydahl, T. Berthing, S. S. Poulsen, O. Schmid, H. Wolff, U. Vogel, Toxicol. Appl. Pharmacol. 2020, 386, 114830.

[63] P. B. Stathopulos, G. A. Scholz, Y.-M. Hwang, J. A. O. Rumfeldt, J. R. Lepock, E. M. Meiering, Protein Sci. Publ. Protein Soc. 2004, 13, 3017.

[64] R. Mawson, M. Rout, G. Ripoll, P. Swiergon, T. Singh, K. Knoerzer, P. Juliano, Ultrason. Sonochem. 2014, 21, 2122.

[65] P. Umek, P. Cevc, A. Jesih, A. Gloter, C. P. Ewels, D. Arčon, Chem. Mater. 2005, 17, 5945.

[66] P. Umek, R. C. Korošec, B. Jančar, R. Dominko, D. Arčon, J. Nanosci. Nanotechnol. 2007, 7, 3502.

[67] P. Umek, C. Bittencourt, P. Guttmann, A. Gloter, S.D. Škapin, D. Arčon, J. Phys. Chem. C 2014, 118, 21250.

