## Supporting Information for "Controlled Fluorescent Labelling of Metal Oxide Nanoparticles for Artefact-free Live Cell Microscopy"

Raw data are available upon request

### Overview of the Labelling Process and Checkpoints to Control it

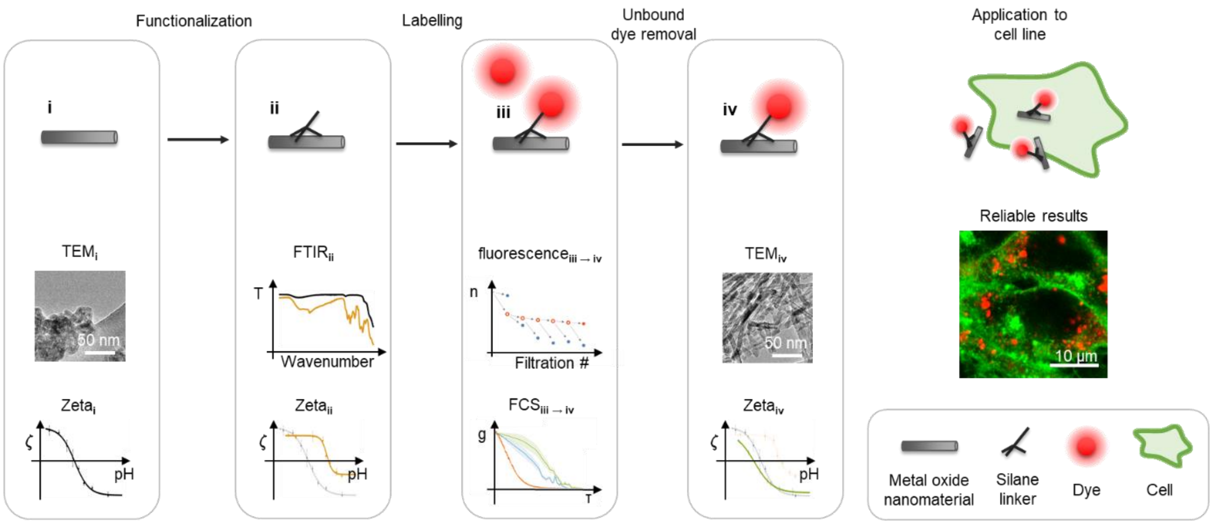

*Figure S1: A schematic of the checkpoints during the labelling procedure, with an emphasis on important checkpoints with regards to the stages of labelling instead of different experimental outcomes.*

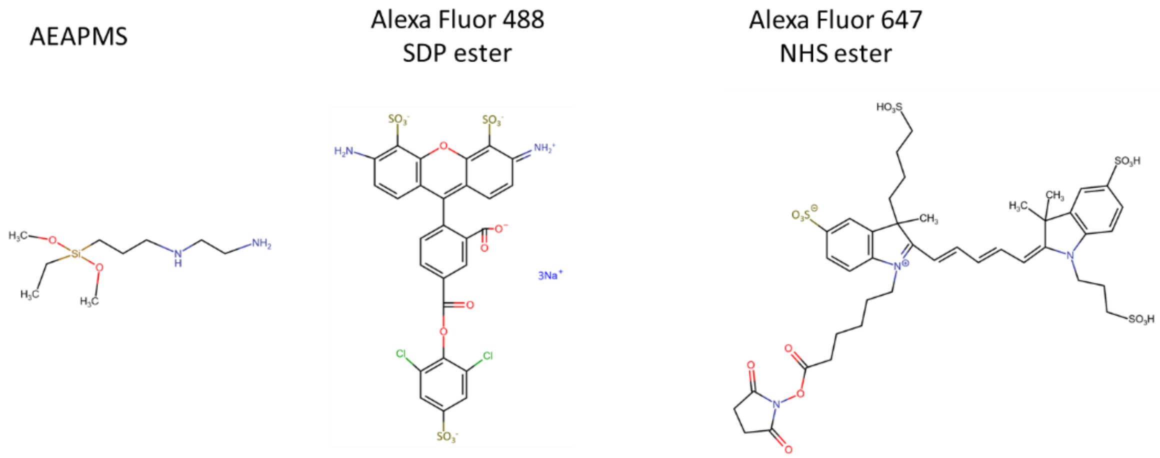

*Figure S2: Structural formulas of AEPMS (3-(2-aminoethylamino)propyltrimethoxysilane, M = 222.36 g/mol) silane linker, the Alexa Fluor 488 SDP ester (M = 825 g/mol), and the Alexa Fluor 647 NHS ester dye (M = 1250 g/mol).*

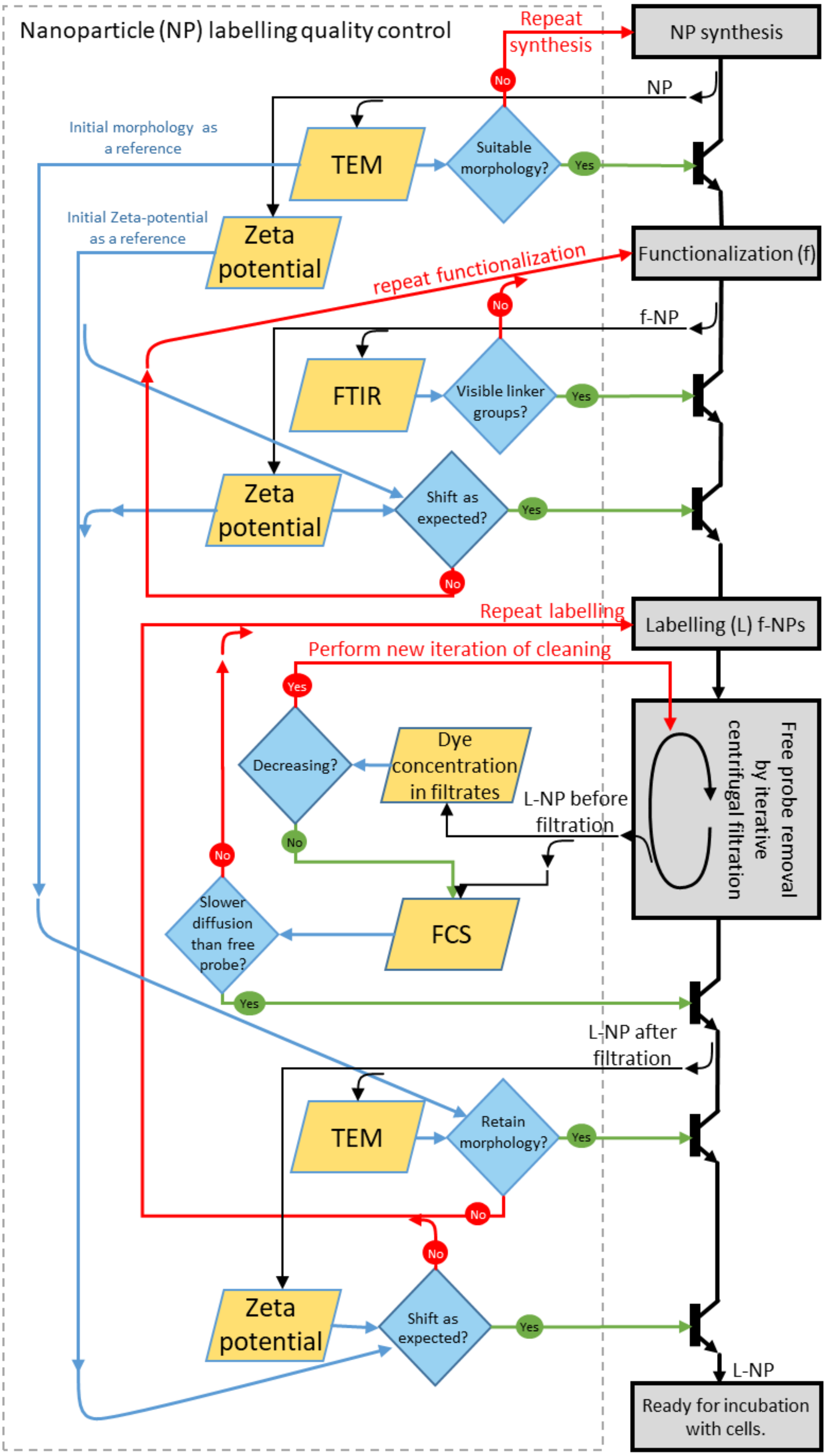

*Figure S3: A flowchart of quality checkpoints during the labelling procedure. NP stands for nanoparticles, f-NP for functionalized nanoparticles and L-NP for labelled nanoparticles.*

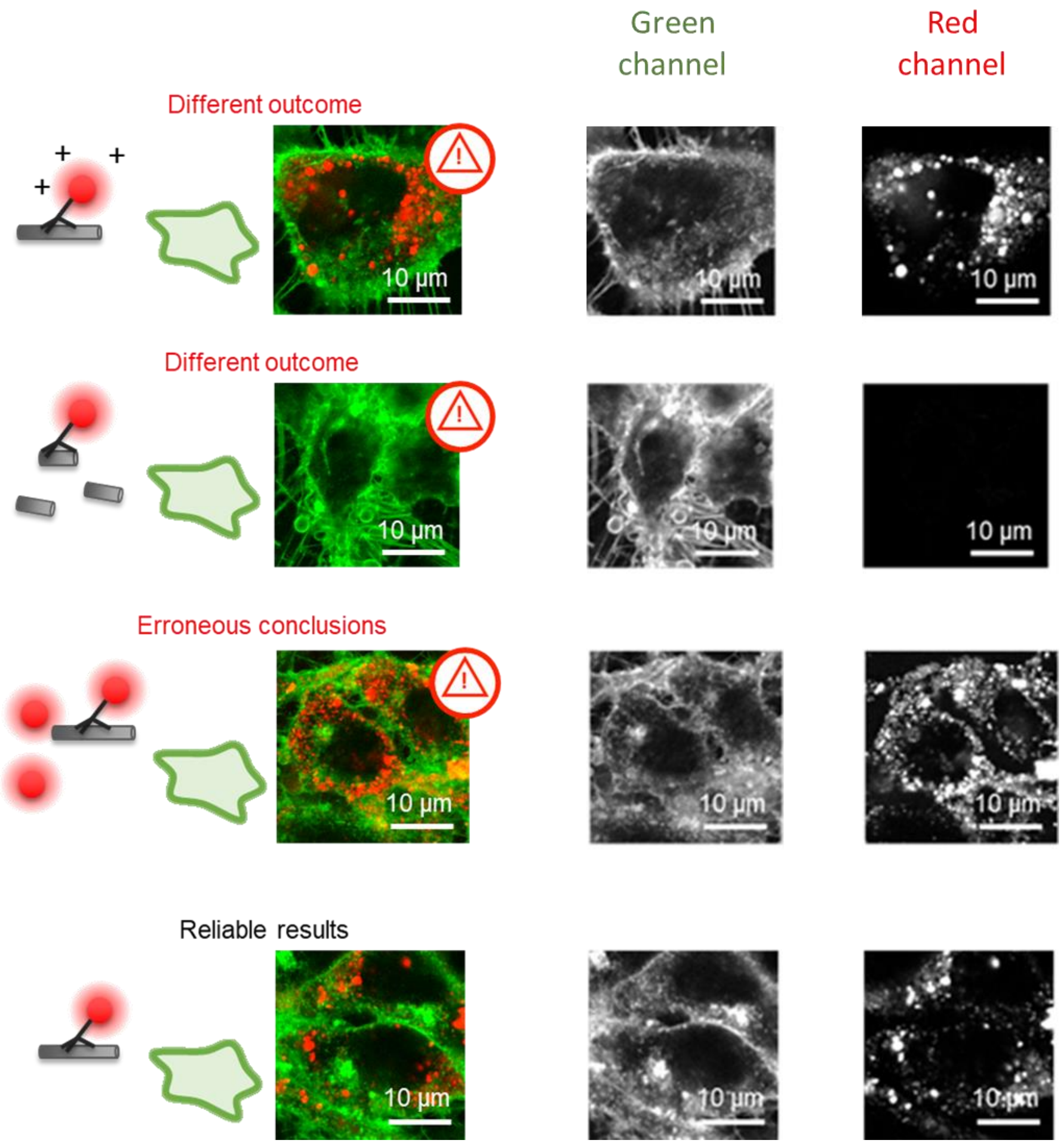

*Figure S4:Separate channels of micrographs from Figure 1b (from the main text) where LA-4 cells (membranes labelled with CellMask Orange, shown in green) were incubated with the exemplary nanomaterial – $TiO_2$ nanotubes – that were either adequately (bottom-most) or inadequately (top three images) fluorescently labelled with Alexa Fluor 647 (shown in red).*

### FTIR Measurements

To show that the surface of $TiO_2$ nanotubes was successfully modified with the silane linker (AEAPMS), ATR FTIR spectra of native $TiO_2$ nanotubes, silane linker and functionalized $TiO_2$ nanotubes are compared (structural formulas of AEAPMS and solvents are shown in Figure S5 and the ATR FTIR spectra is shown in Figure S6). The ATR FTIR spectrum of $TiO_2$ nanotubes shows a main broad peak starting below 1000 $cm^{-1}$ which is attributed to Ti-O stretching and Ti-O-Ti bridging stretching modes.[1,2] Beside this peak two low intensity peaks are observed; a broad one centered at 3329 $cm^{-1}$ and the second one at 1635 $cm^{-1}$ belonging to O-H stretching and H-O-H bending modes, respectively. From this it can be concluded that surface of $TiO_2$ nanotubes is covered with a thin water layer. In the spectrum of AEAPMS there are several characteristic bands; (i) N-H stretching at around 3300 $cm^{-1}$ (in -NH- and -$NH_2$ groups), (ii) C-H stretching at 2941 $cm^{-1}$ (in -$CH_2$- groups), 2844 $cm^{-1}$ (in Si-O-$CH_3$ group) and a sharp intensive band at 1079 $cm^{-1}$ (in Si-O-$CH_3$ group). In the spectra of functionalized NTs in comparison to native $TiO_2$ nanotubes several new bands appear: (i) at 2931 and 2874 $cm^{-1}$ characteristic to C-H stretching vibrations, (ii) 1029 $cm^{-1}$ corresponding to Si-O-Si vibration, and (iii) a weak peak at 922 $cm^{-1}$ due to Ti-O-Si vibration [1,3] which provide important information that the silane linker was successfully grafted at the surface of $TiO_2$ nanotubes.

[1] J. Zhao, M. Milanova, M. M. C. G. Warmoeskerken, V. Dutschk, *Colloids Surf. Physicochem. Eng. Asp.* **2012**, *413*, 273.

[2] A. M. Peiró, J. Peral, C. Domingo, X. Domènech, J. A. Ayllón, *Chem. Mater.* **2001**, *13*, 2567.

[3] P. Launer, B. Arkles, in *Silicon Compd. Silanes Silicones*, Gelest Inc., Morrisville, PA, USA, **2013**.

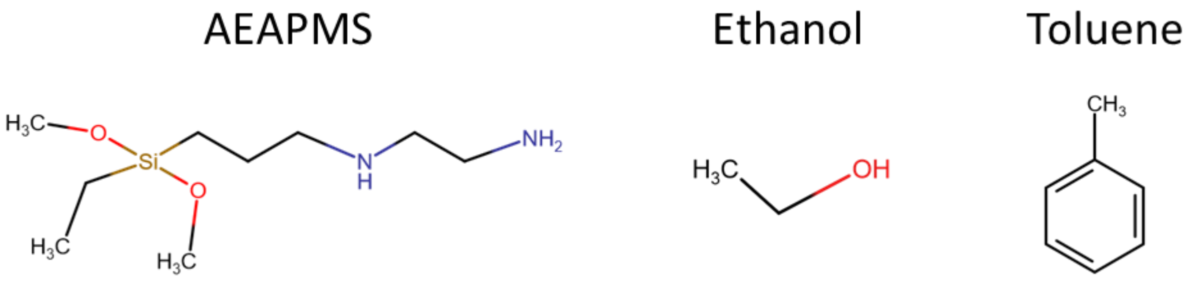

*Figure S5: Structural formulas of the silane linker AEAPMS (3-(2-aminoethylamino)propyltrimethoxysilane) and both solvents used during functionalization - ethanol and toluene. Note that the C-N, N-H, Si-O, and Si-C bonds are present only in AEAPMS.*

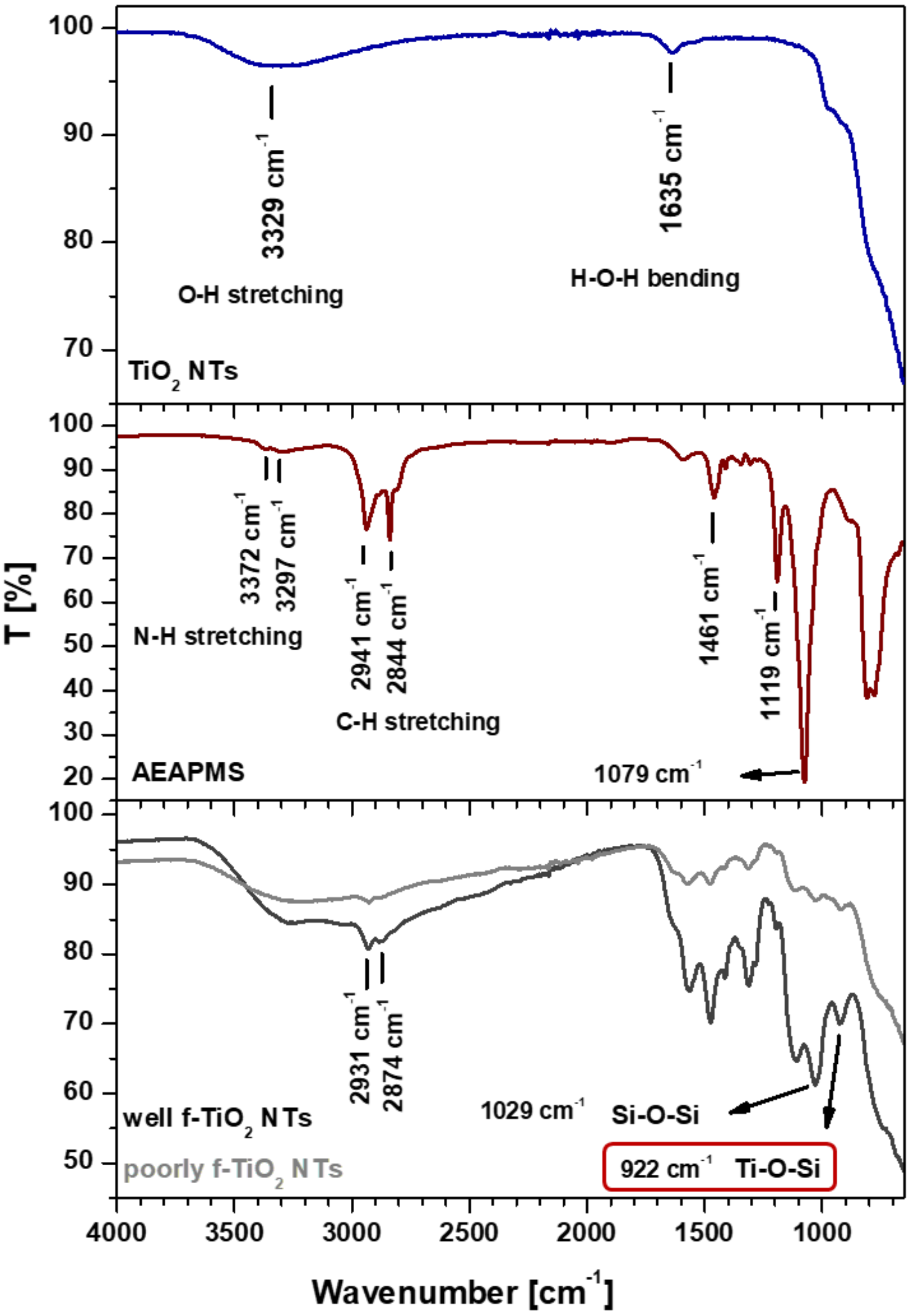

*Figure S6: Comparison of ATR FTIR spectra, from top to bottom, of $TiO_2$ nanotubes, silane linker (AEAPMS; 3-(2-aminoethylamino)propyltrimethoxysilane) and well and poorly functionalized $TiO_2$ nanotubes (well and poorly f-$TiO_2$ NTs). The newly-formed bond between the linker and the $TiO_2$ surface is highlighted in red.*

### Zeta Potential Measurements

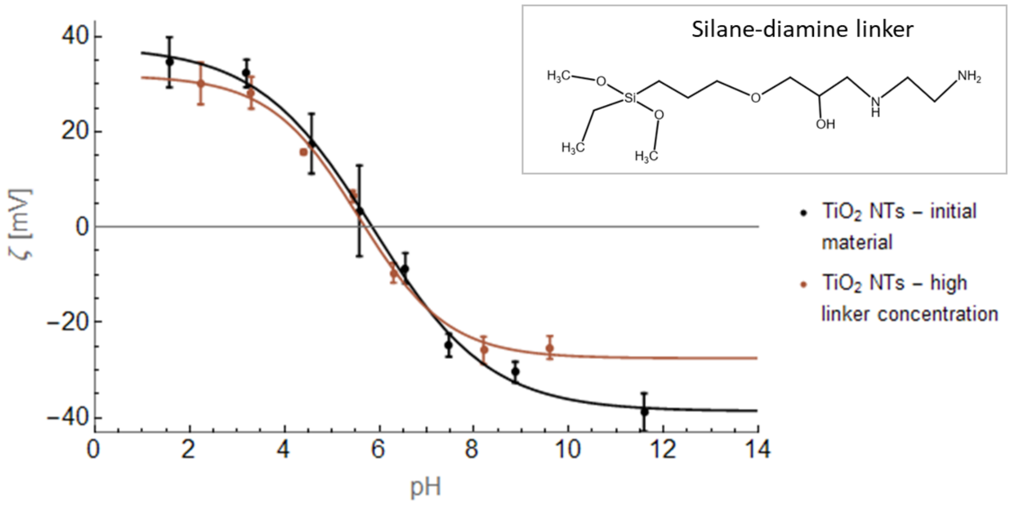

*Figure S7: An example of a Zeta potential measurement of $TiO_2$ nanotubes, unsuccessfully functionalized using a silane and diamine linker as an alternative to AEAPMS (as shown in Figure S8). The Zeta potential of correctly functionalized nanomaterial would move towards the right (similar as the one shown in the main text). However, the Zeta potential of the samples with large quantities of linker (brown) is approximately the same as that of the native $TiO_2$ nanotubes (black), suggesting that the silane linker in the sample is only partially bound to the nanomaterial (with one instead of three bounds), and the reaction between the diamine and the silane linker did not proceed correctly.*

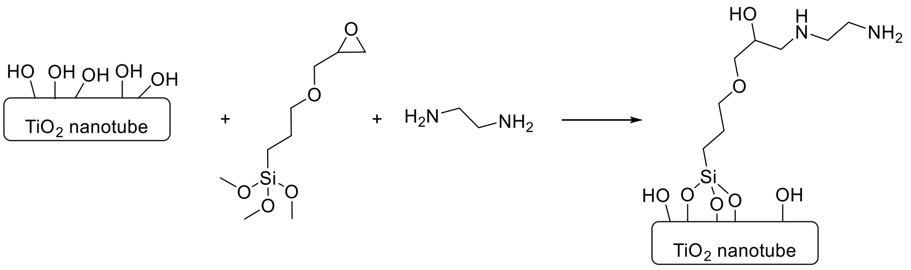

*Figure S8: A schematic of a functionalization process, resulting in the Zeta potential measurements shown in Figure S7. $TiO_2$ nanotubes were suspended in water, ethanol and 0.1 M NaOH were added to achieve a pH of 11. Then the silane linker end ethylene diamine were added and the reaction mixture was stirred overnight at 50 ˚C.*

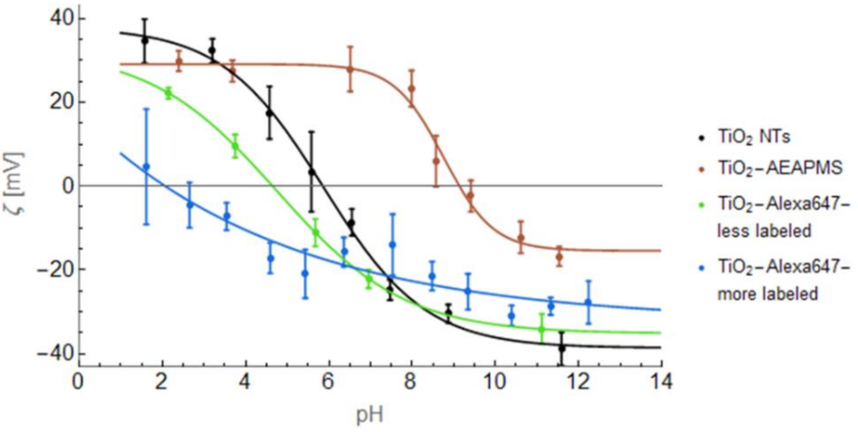

*Figure S9: Zeta potential measurements of native $TiO_2$ (black), functionalized $TiO_2$-AEAPMS (brown), $TiO_2$ labelled with copious amount of Alexa647 (blue) and labelled with just enough Alexa647 to bring the Zeta potential back to the native one (green).The less labelled $TiO_2$-Alexa647 was still visible under the fluorescent microscope, but not as brightly as the more labelled $TiO_2$-Alexa647.*

### Selection of Fluorescent Dye

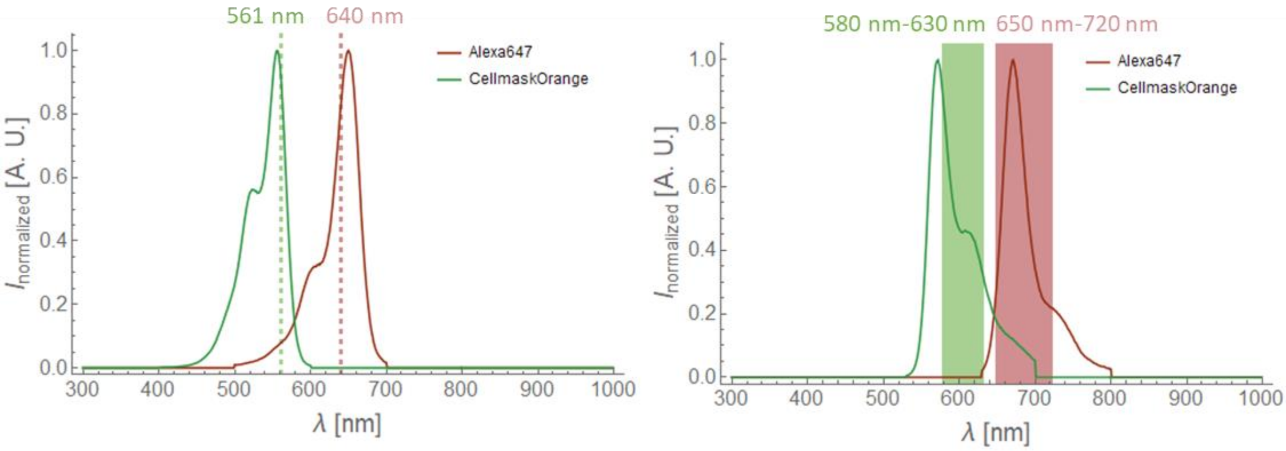

*Figure S10: Selected dyes used in fluorescent microscopy experiments (Alexa Fluor 647, red, and CellMask Orange, green) with corresponding excitation lasers and filters sets.*

### Filtration – Removal of Free Dye

Filtrates of initial sample and retentates and filtrates of consecutive filtrations are presented in Figure S11 and Figure S12 to visually demonstrate the decreased dye concentrations after filtration. The same Figure is also used to stress out the importance of sonication of centrifugal filter device (CFD).

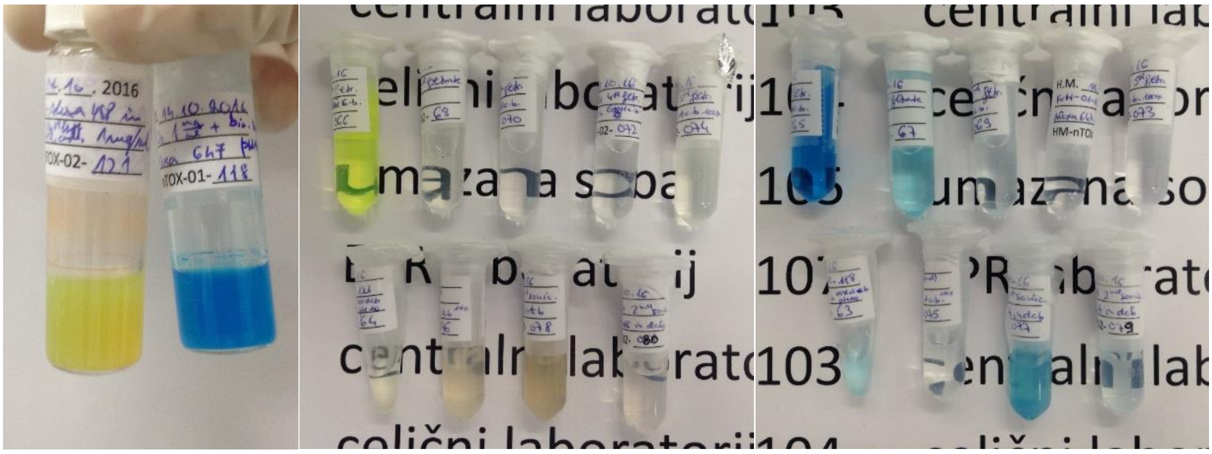

*Figure S11: Filtration of $TiO_2$-Alexa488 (yellow, middle) and $TiO_2$-Alexa647 (blue, right). Left: the freshly labelled nanomaterial just prior to the first filtration. Note the turbidity due to presence of nanomaterial (1 mg/mL), and strong color due to a large concentration of (unbound) fluorescent dye. Filtrates and retentates of the filtration of the $TiO_2$-Alexa488 (middle) and $TiO_2$-Alexa647 (right) shown on the left: the samples in the upper rows are the $1^{st}$ to $5^{th}$ filtrates (from left to right) and samples in the lower rows are (from left to right): 10-times diluted starting sample, retentate after the $5^{th}$ filtration pipetted directly from the centrifugal filter device (CFD), the retentate after the $1^{st}$ sonication of CFD, and the retentate after $2^{nd}$ sonication of CFD.*

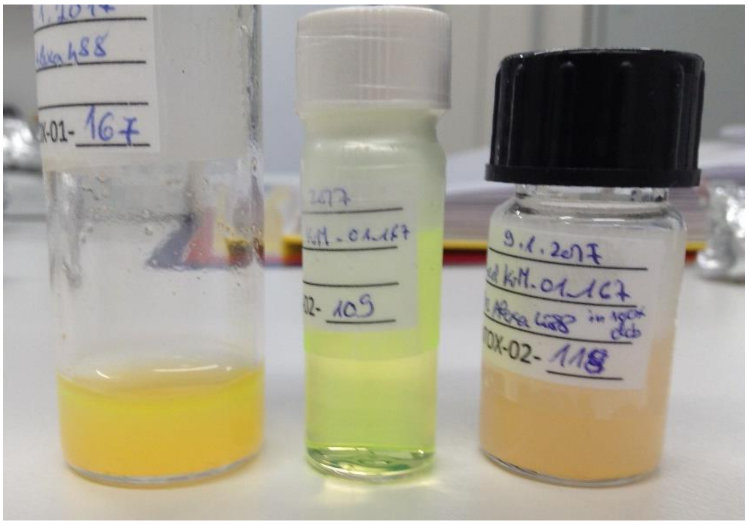

*Figure S12: Filtration of 1 mg/mL $TiO_2$ nanotubes, labelled with Alexa Fluor 488: labelled nanomaterial prior to first filtration, first filtrate and the well-filtered sample (from left to right, undiluted samples).*

### Measuring Fluorescence Intensity to Evaluate the Filtration

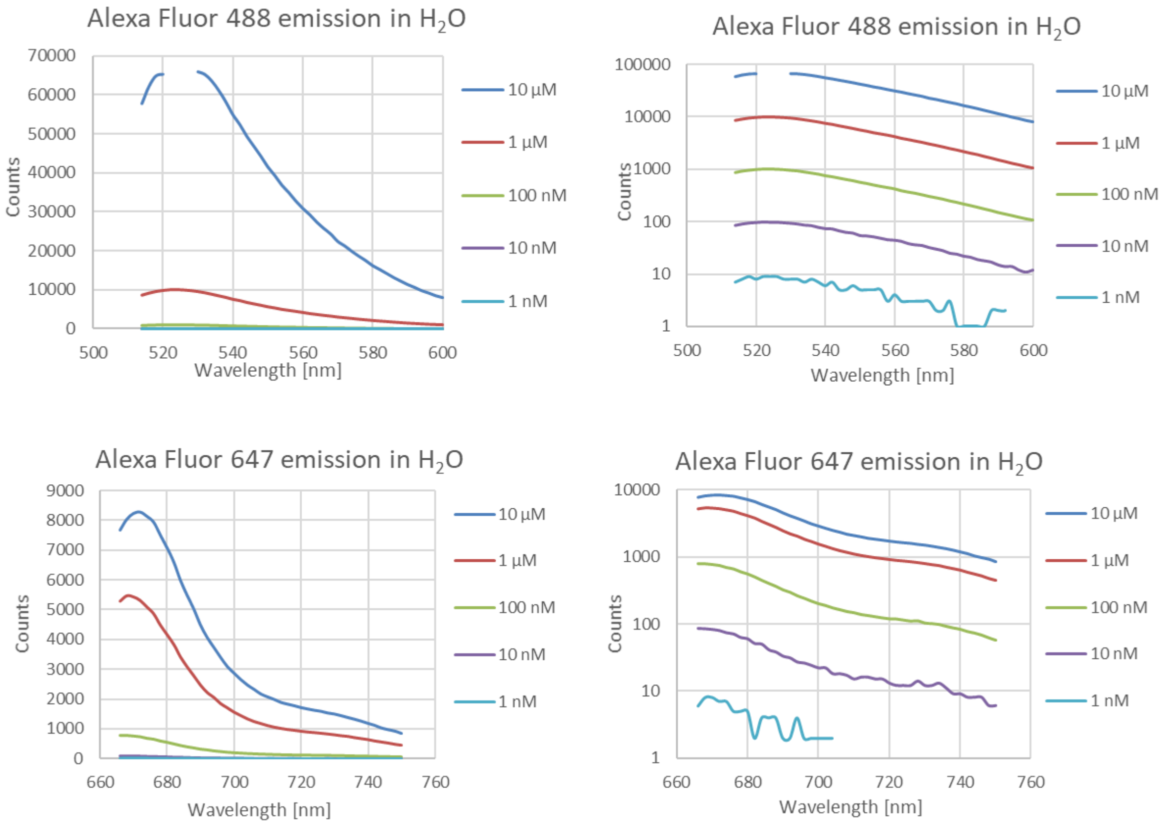

*Figure S13: Emission spectra of Alexa Fluor 488 (above) and Alexa Fluor 647 (below) shown in linear (left) and logarithmic scale (right). The emission was measured with a spectrophotometer at different concentrations of the dyes in $H_2O$ (as shown in the legends) and the background spectrum was subtracted from the measurement. Measurements of Alexa Fluor 488 at 10 µM (blue curve on upper graphs) are missing some measurements at the peak intensity due to saturation of the detector at the set gain of the detector.*

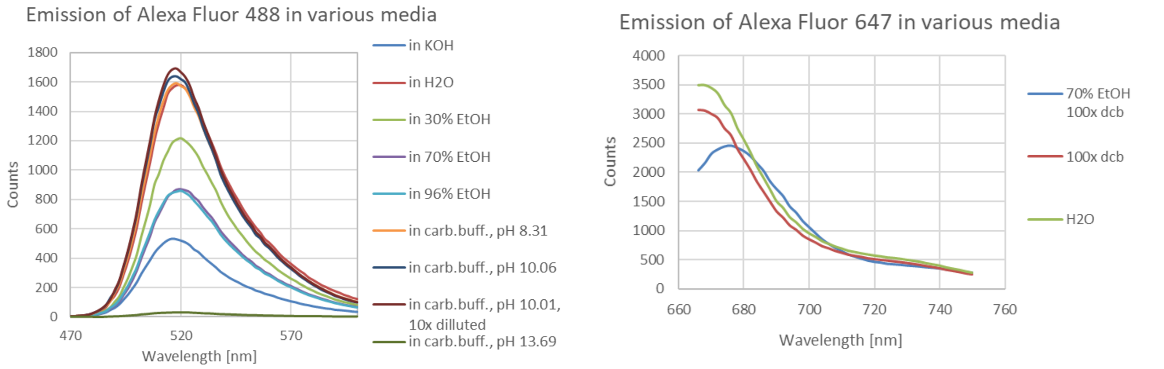

*Figure S14: Influence of various media on the emission spectra of Alexa Fluor 488 (left) and Alexa Fluor 647 (right). The background spectrum was subtracted from the measurement.*

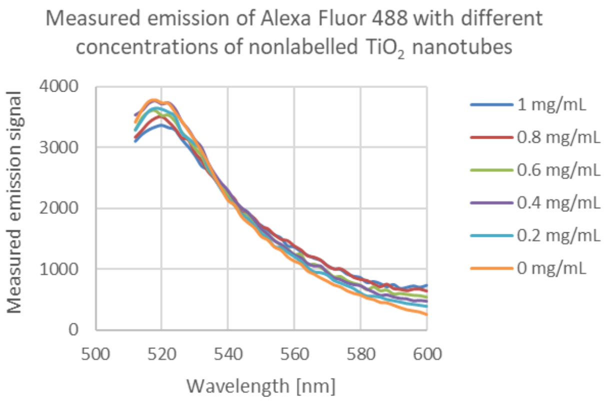

*Figure S15: Influence of the scattering of light from $TiO_2$ nanotubes on the measured emission spectrum of the sample. Free Alexa Fluor 488 was mixed with various concentrations of nonlabelled $TiO_2$ nanotubes to induce scattering. Measurements were performed in 0.1 M KOH to prevent nanomaterial aggregation. The background spectrum was subtracted from the measurements.*

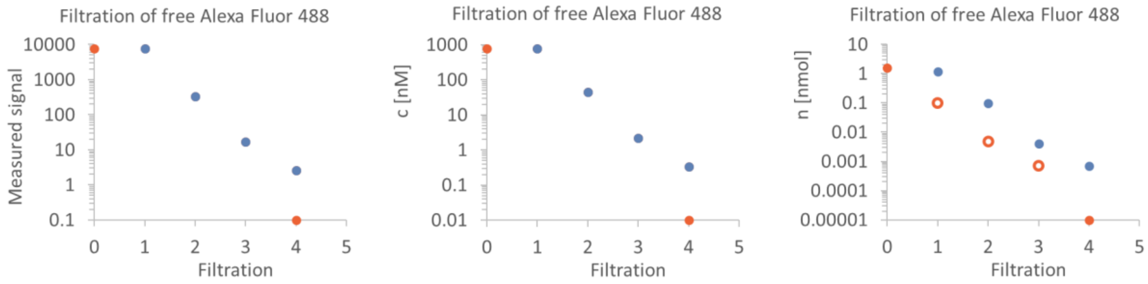

*Figure S16: Exemplary calculation of amount of dye in filtrates (blue) and retentates (orange) from intensities measured with a spectrophotometer. The empty orange markers were calculated as the difference between subsequent filtrates. The left plot is a read-out of the maximum of the measured fluorescence emission spectrum, which was measured exactly the same as the calibrations in Figure S13, Figure S14, and Figure S15 (same sample volumes, excitation wavelength, and measurement settings: gain, bandwidth, number of flashes, flash frequency, integration time and z position). The middle plot was calculated from the measured signal and corrected for dilution, medium and nanomaterial scattering. From that and the volumes of the samples, the values for the right plot were calculated.*

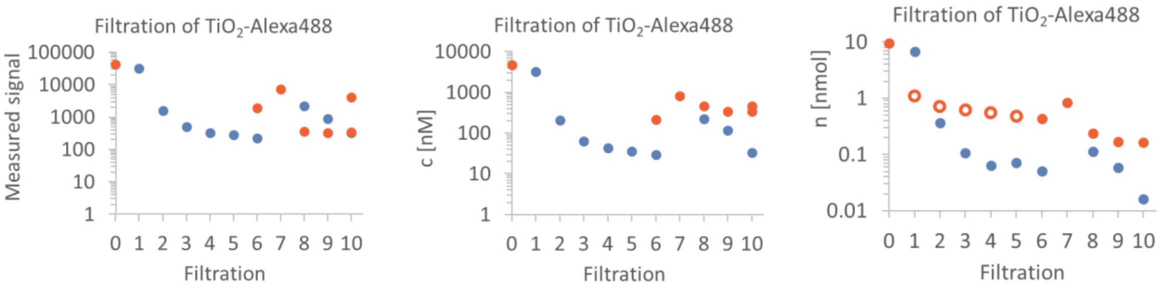

*Figure S17: Example of calculation of amount of dye in filtrates (blue) and retentates (orange) from measured signal. The empty orange markers were calculated as the difference between measured samples. The left plot is a read-out of the maximum of the measured fluorescence emission spectrum, which was measured exactly the same as the calibrations in Figure S13, Figure S14, and Figure S15 (same sample volumes, excitation wavelength, and measurement settings: gain, bandwidth, number of flashes, flash frequency, integration time and z position). The middle plot was calculated from the measured signal and corrected for dilution, medium and nanomaterial scattering. From that and the volumes of the samples, the values for the right plot were calculated.*

### Estimation of the Number of Fluorophores per Nanotube

We measured that a 1 mg/mL sample of our labelled nanomaterial has the same signal as 300 nM concentration of free fluorescent dye, which we estimated to correspond to 1.5 fluorophores per nanotube, as described in detail below.

For this assessment, one must estimate:

i) the number of nanoparticles in 1 mg of material
ii) the number of dyes in 1 mg of material (via the measured fluorescence intensity)

i) From TEM images of the nanomaterial, we estimated that our nanotubes are hollow tubes with a mean length $L = 200$ nm and a diameter $D = 2R = 10$ nm. Because they are hollow, only their outer surface is available for linker and dye attachment. One can calculate that the outer surface of a nanotube is approximately $S_{NT} = 2\pi RL = 6 \cdot 10^{-15}$ m$^2$.

The measured Brunauer–Emmett–Teller (BET) surface of the nanotubes is $BET = 150$ m$^2$/g, with half of this value corresponding to the outer surface. The number of nanotubes in one milligram of material is thus estimated to be

$$\frac{\#_{NT}}{1\text{ mg}} = \frac{0.5\ BET}{S_{NT}} = \frac{75\text{ m}^2/\text{g}}{6 \cdot 10^{-15}\text{ m}^2} = 12 \cdot 10^{13}\text{ nanotubes/mg}$$

ii) According to the calibration curves in the previous chapter, the labelled nanotubes with a concentration of 1 mg/mL have the same signal as 300 nM free dye. Hence, the number of dye molecules in 1 mg of labelled nanomaterial is

$$\frac{\#_{dye}}{1\text{ mg of nanotubes}} = \frac{300\text{ nM}}{1\text{ mg/mL}} = \frac{300 \cdot 10^{-9}\text{ mol/L}}{1\text{ mg/mL}} = \frac{300 \cdot 10^{-9}\ \ 6 \cdot 10^{23}\text{dyes/L}}{1\text{ mg/mL}} =$$

$$= 18 \cdot 10^{13}\text{dyes/mg of nanotubes}$$

Consequently, in this case we had $18 \cdot 10^{13}$ dyes per $12 \cdot 10^{13}$ nanotubes, corresponding to 1.5 dyes per nanotube. As it turned out from other experiments, described in the main text, this degree of labelling still enabled imaging with our setup, but did not majorly influence the nanomaterial surface.

### Cellular Distribution and Signal of Free Dye

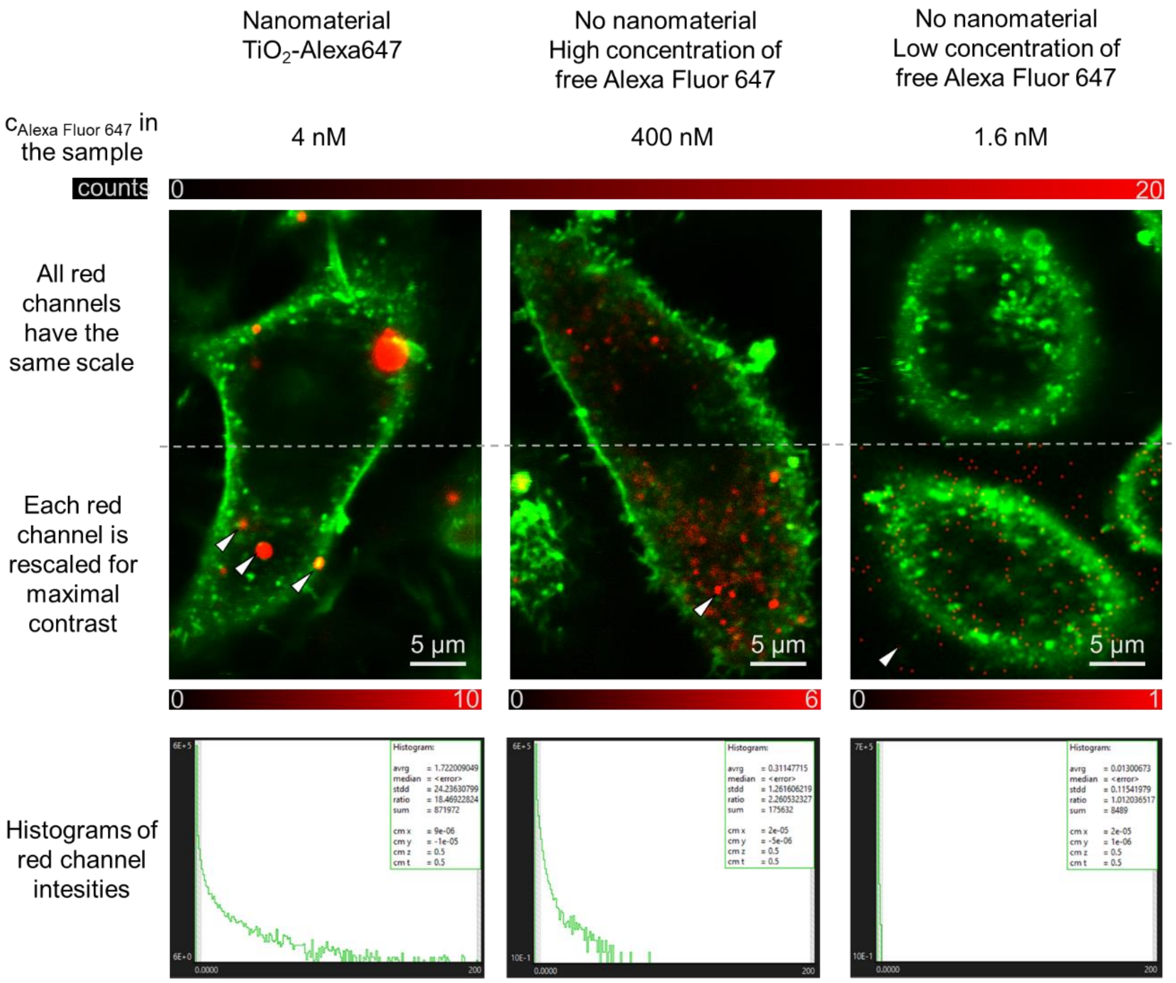

*Figure S18: A comparison of LA-4 cells (labelled with CellMask Orange, shown in green), incubated for two days with (left to right) a 1:1 ratio between surface of nanomaterial and surface of cells (surface dose) of $TiO_2$-Alexa647 (Alexa Fluor 647 signal shown in red, concentration of bound Alexa Fluor 647 in the sample is approx. 4 nM), a high concentration of free Alexa Fluor 647 (400 nM), and a low concentration of free Alexa Fluor 647 (1.6 nM). The red channels in the upper half of the micrographs were all contrasted from 0 to 20 counts to ease the comparison of the signals. In the lower half, the red channels were contrasted differently for each sample in order to show the distribution of Alexa Fluor 647 signal in the cell. The arrows indicate the red signal present in the cell and on the membrane (left), inside the cell in the vesicles (middle) and noise only (right). In the bottom row equally scaled distributions of Alexa Fluor 647 signal in the images are presented for comparison. Note that even if all dye desorbed from the well-labelled nanoparticles at a 1:1 surface dose of nanomaterial, the final dye concentration in the sample would be 4 nM and would hence be negligible. For micrographs of separate color channels refer to Figure S19.*

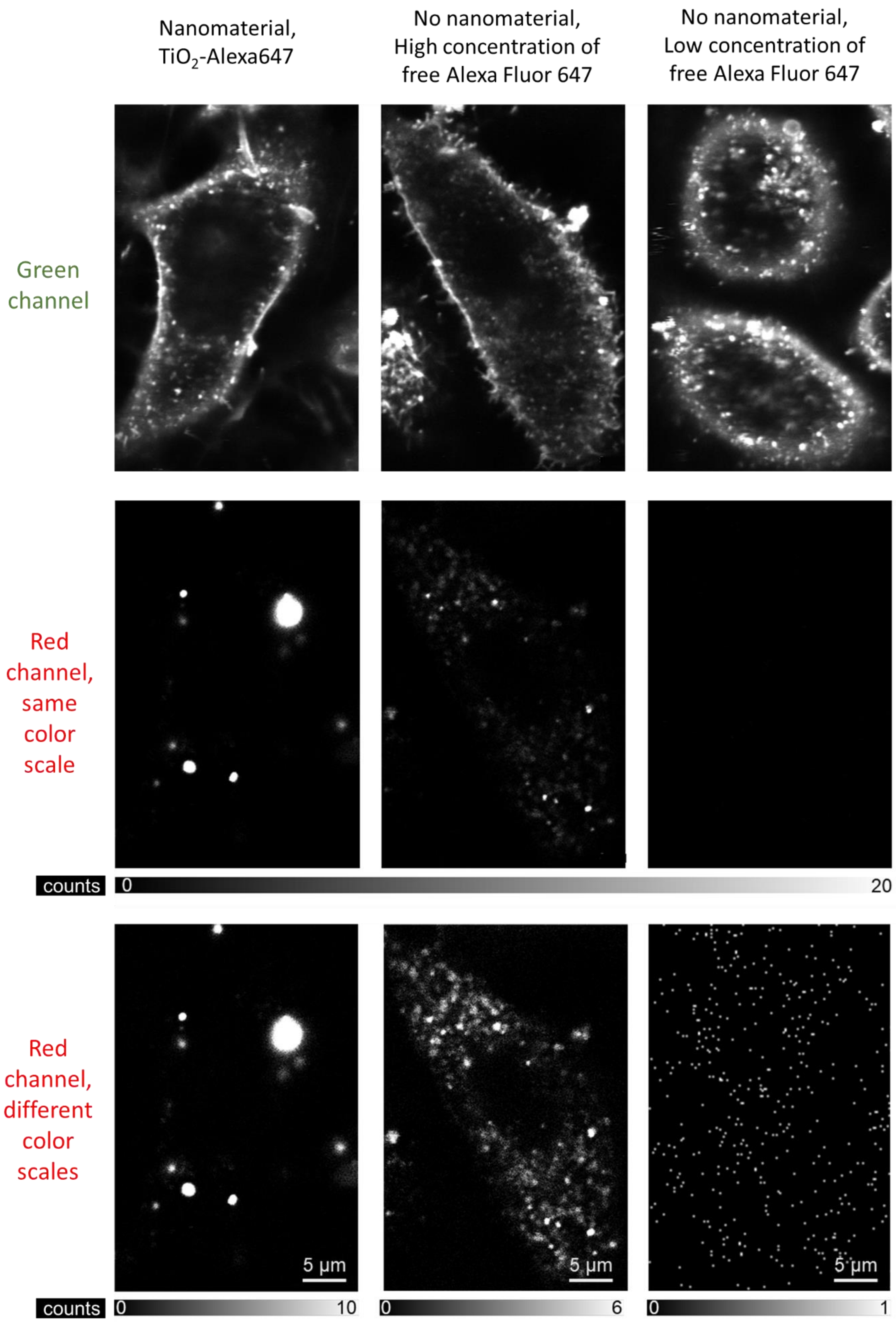

*Figure S19: Separate channels of micrographs from Figure S18 showing a comparison of LA-4 cells (labelled with CellMask Orange, green channel), incubated for two days with (left to right) a 1:1 ratio between surface of nanomaterial and surface of cells (surface dose) of $TiO_2$-Alexa647 (Alexa Fluor 647 in red channel, concentration of bound Alexa Fluor 647 in the sample is approx. 4 nM), a high concentration of free Alexa Fluor 647 (400 nM), and a low concentration of free Alexa Fluor 647 (1.6 nM).*

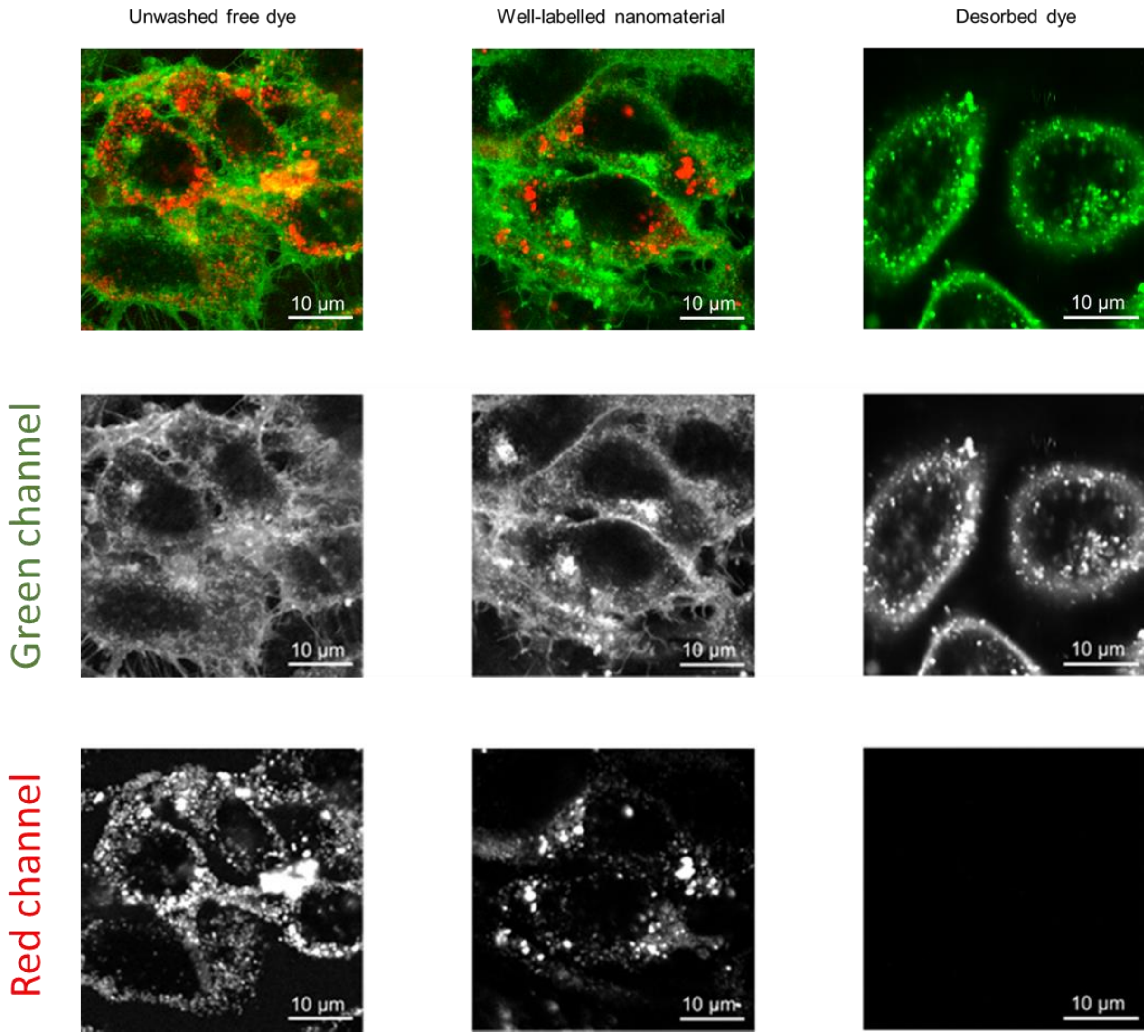

*Figure S20: Separate channels of micrographs from Figure 3d (from the main text) showing LA-4 cells (cell membranes labelled with CellMask Orange, shown in green) exposed to labelled $TiO_2$ nanotubes with various amounts of free dye in the sample (the bound and unbound dye is Alexa Fluor 647, shown in red and scaled so its fluorescence intensity is directly comparable across the images).*

### FCS Measurements

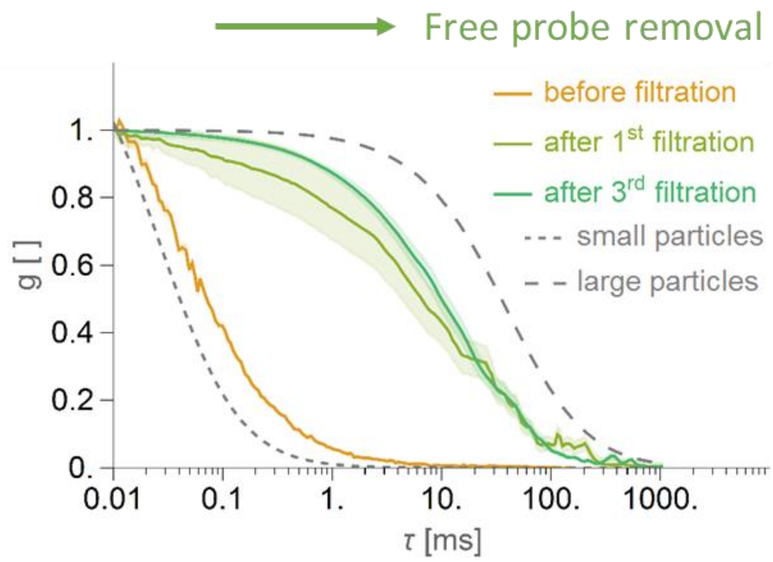

*Figure S21: Monitoring of filtration of labelled nanoparticles using FCS: with each passing filtration, the retentate should contain a larger fraction of slow-moving particles, which reflects in autocorrelations shifted towards longer lag times (τ). When the difference between curves of retentates from subsequent filtrations is negligible, the free dye has been successfully removed.*

### Sonication of Nanomaterial

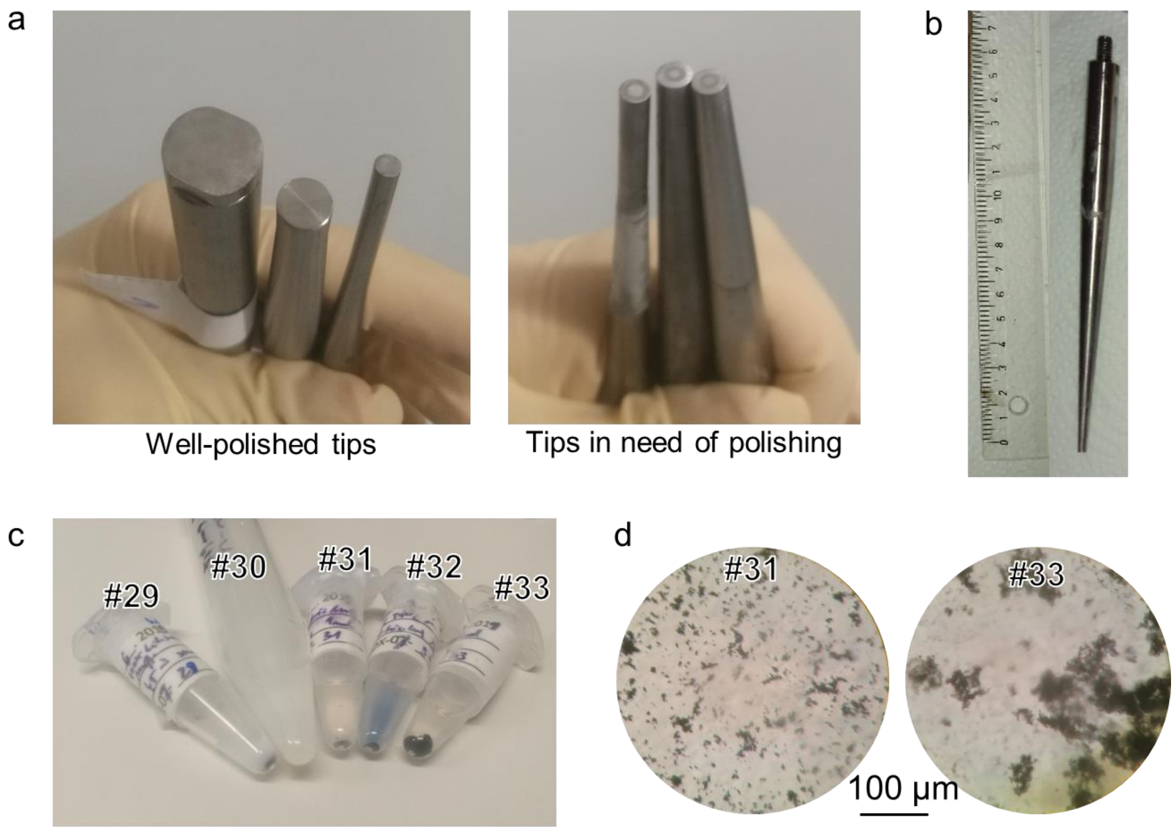

*Figure S22: a) An example of well-polished tips (left) and tips after sonication, exhibiting characteristic dark circles, indicating that they should be polished prior to the next sonication (right). b) The 3-mm (30 mm$^2$) tip used for most of our sonications. c) Five samples in preparation of which different degrees of polishing and cleaning of the tip prior to sonication were applied, resulting in different amounts of tip debris in the sonicated sample: from none (#30) to some (#29, #31, #32), and a lot (#33). Different conditions are described in Table S1. The tip debris can be observed as black sediment on the bottom of the Eppendorf tubes. Note that this is certainly not nanomaterial, as the nanomaterial used forms a milky white suspension (as #30; note that the clear samples #31 and #33 did not contain any nanomaterial at all). d) Micrographs of sonicated media without nanomaterial contain large amounts of tip debris (black structures).*

*Table S1: Description of samples shown in Figure S22 C and D. The washing of the tip in sample #30 consisted of thoroughly washing the tip under running water and a 1-minute long sonication in ethanol followed by a quick rinse with DI $H_2O$.*

| *Sample* | *$TiO_2$ concentration* | *Volume of medium* | *Time of sonication* | *Tip polished and washed* | *Tip debris* |
|---|---|---|---|---|---|
| #29 | 1 mg/mL | 1 mL | 45 min | poorly | some |
| #30 | 1 mg/mL | 5 mL | 4 h | yes | none |
| #31 | - | 1 mL | 2 h | poorly | some |
| #32 | 1 mg/mL | 1 mL | 2 h | poorly | some |
| #33 | - | 1 mL | 4 h | poorly | a lot |

### Micrographs of Detected Early Molecular Events

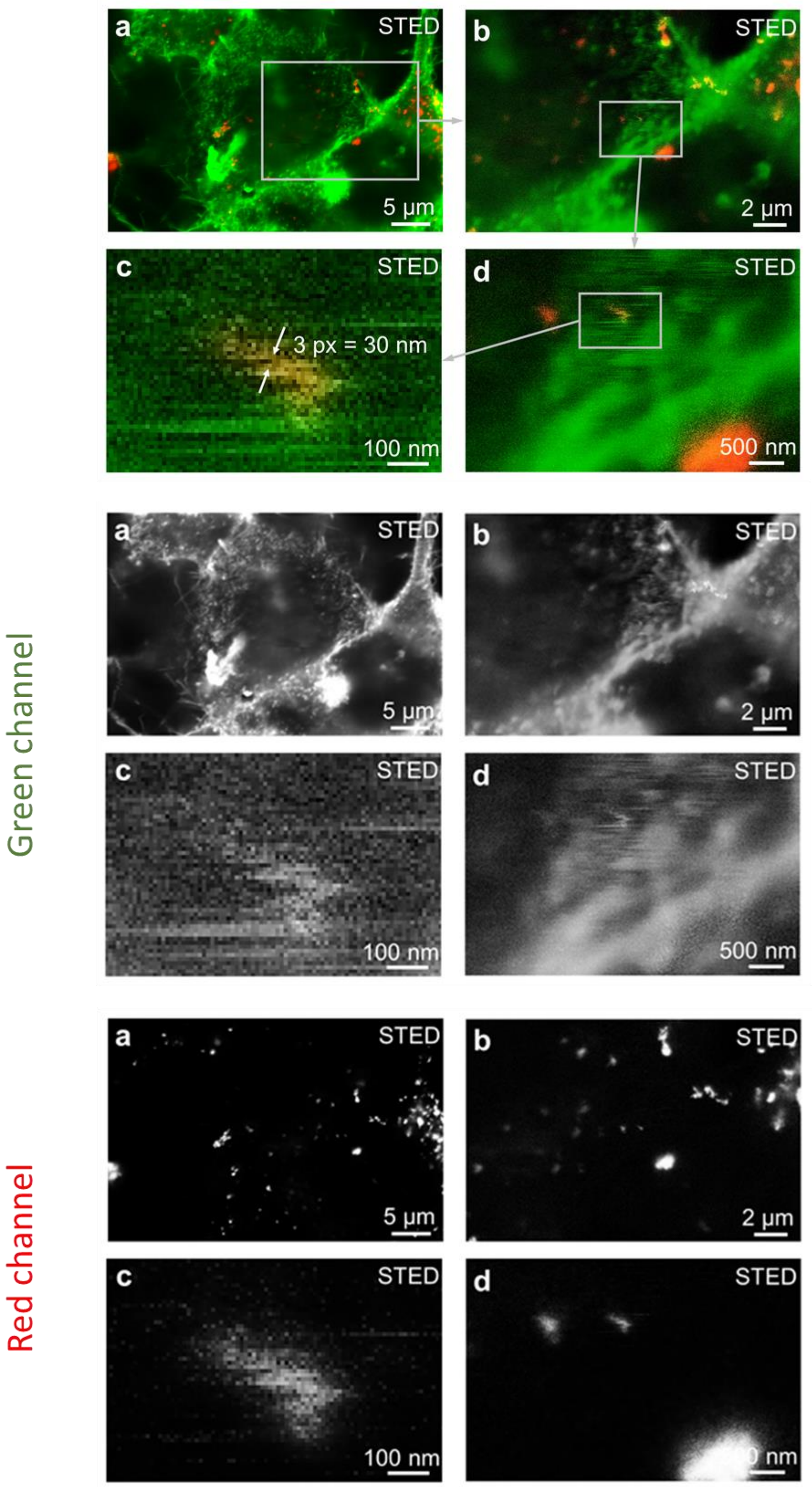

*Figure S23: Separate channels of micrographs from Figure 5 (from the main text) showing LA-4 cells (membranes labelled with CellMask Orange, green) incubated for 2 days with efficiently and stably labelled nanoparticles (Alexa Fluor 647, red). The ratio between the surface of the nanomaterial and surface of cells is 1:1 and the signal of nanomaterial is logarithmically scaled.*

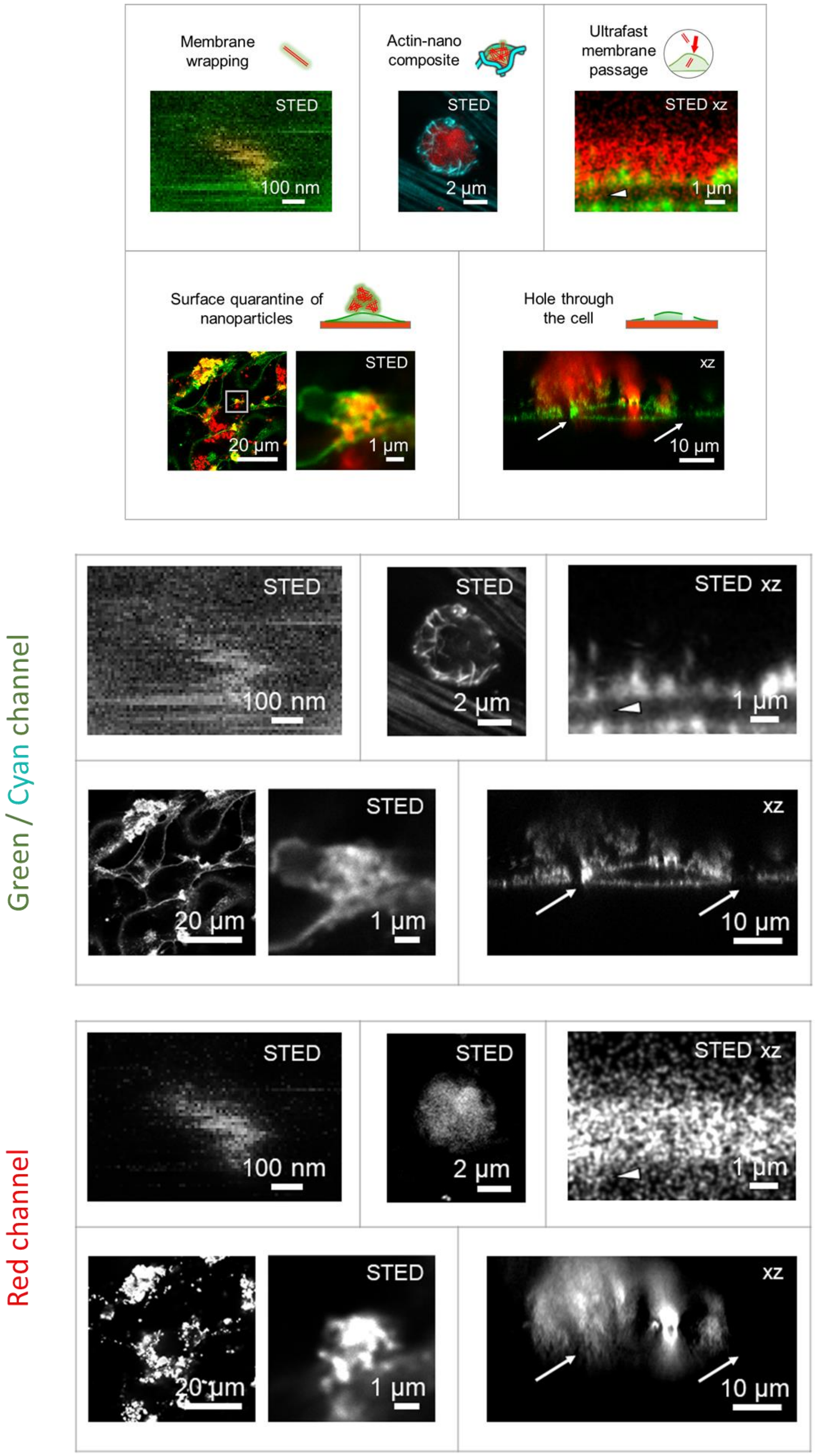

*Figure S24: Separate channels of micrographs from Figure 6 (from the main text) showing five early molecular events in living LA-4 cells following exposure to nanomaterial ($TiO_2$ nanotubes labelled with Alexa Fluor 647, red), detected with confocal or STED fluorescence microscopy. Cell membranes are labelled with CellMask Orange (green), actin with SiR Actin (cyan).*

### Additional Micrographs of Artefacts

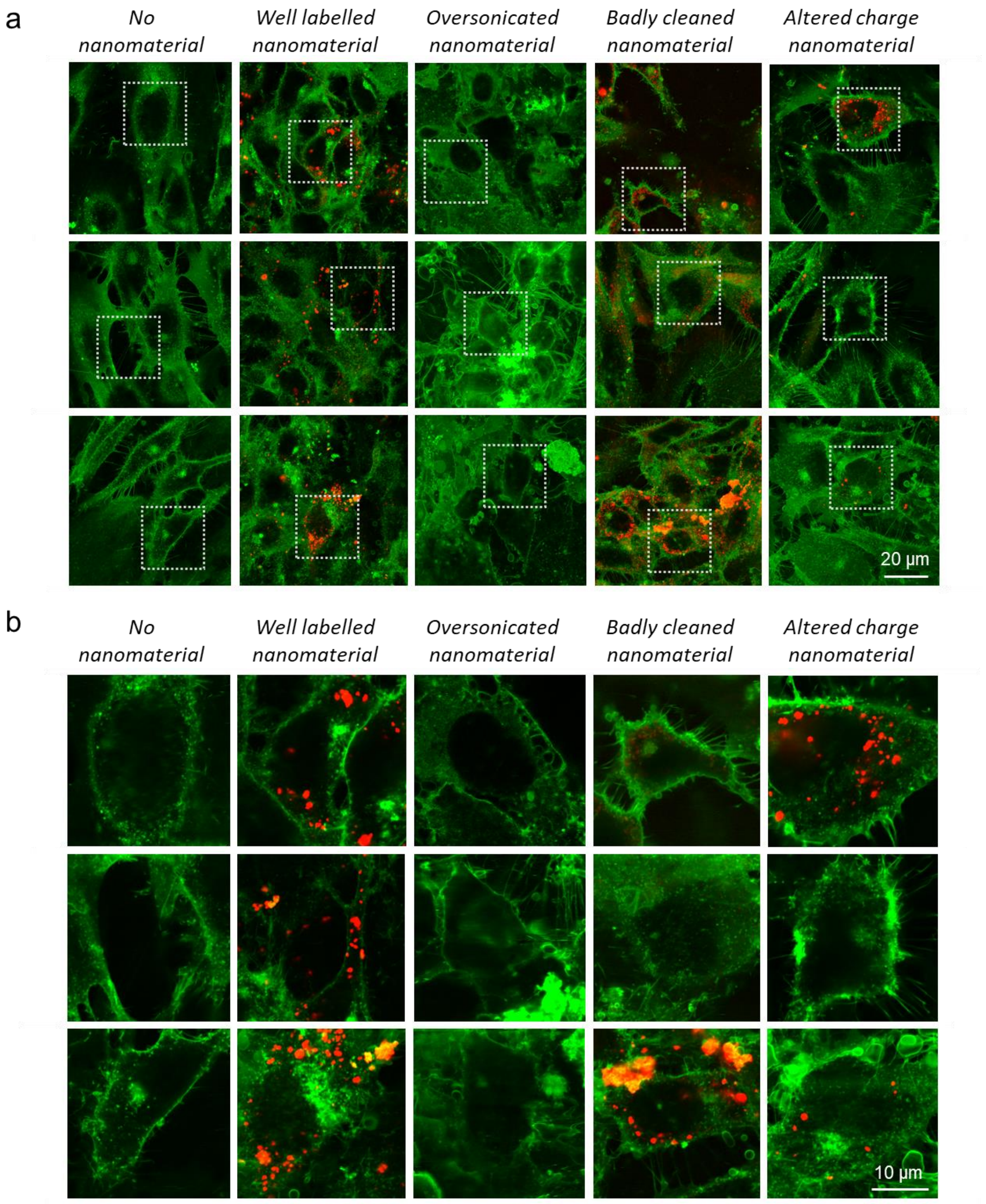

*Figure S25: a) Confocal images and b) corresponding STED zoom-ins of LA-4 cell incubated with $TiO_2$-Alexa647 nanotubes. Membrane is labelled with CellMask Orange (shown in green) and nanoparticles are labelled with Alexa Fluor 647 (shown in red). From left to right: "No nanomaterial" – just the cells at conditions of incubation, "Well labelled nanomaterial" – cells incubated with correctly labelled $TiO_2$-Alexa647 at 1:1 ratio between surface of nanomaterial and surface of cells (surface dose), "Oversonicated nanomaterial" – cells incubated with nanomaterial that was broken in small pieces by extensive sonication, "Badly cleaned nanomaterial" – cells incubated with labelled nanomaterial and a 100-times excess of free dye, "Altered charge nanomaterial" – cells incubated with nanomaterial with altered charge of the surface. Contrast is individually adjusted in each image for best visibility of membrane morphology and nanoparticles distributions. For micrographs of separate color channels refer to Figure S26 and Figure S27.*

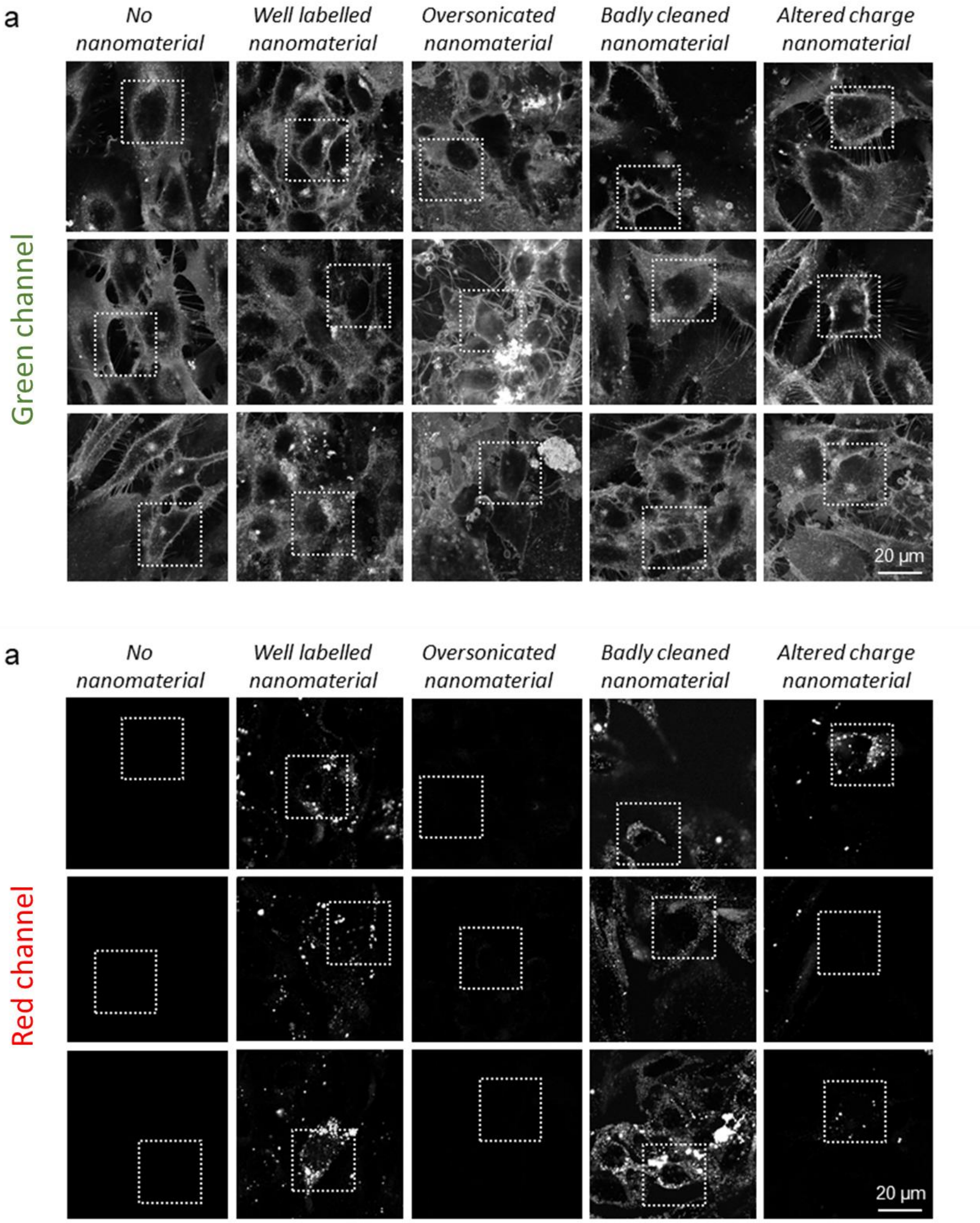

*Figure S26: Separated channels of confocal images shown in Figure S25 – LA-4 cell incubated with $TiO_2$-Alexa647 nanotubes showing various artefacts that can arise due to nanomaterial labelling. The cell membranes (labelled with CellMask Orange), are shown in the green channel, and nanoparticles (labelled with Alexa Fluor 647) in the red channel.*

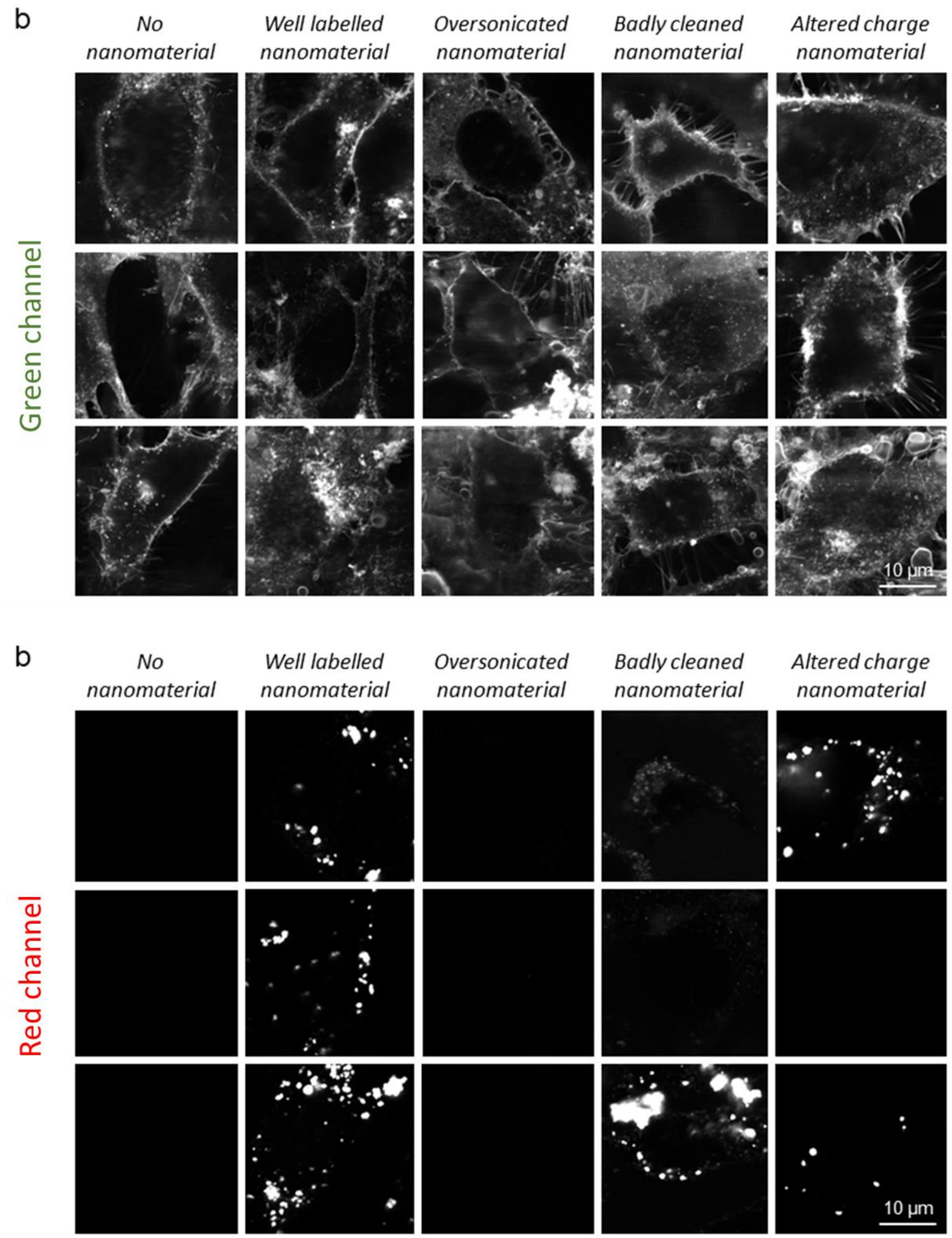

*Figure S27: Separated channels of STED zoom-ins shown in Figure S25 – LA-4 cell incubated with $TiO_2$-Alexa647 nanotubes showing various artefacts that can arise due to nanomaterial labelling. The cell membranes (labelled with CellMask Orange), are shown in the green channel, and nanoparticles (labelled with Alexa Fluor 647) in the red channel.*

### A Detailed Procedure for Labelling $TiO_2$ Nanotubes with Alexa Fluor 488 SDP Ester Using AEAPMS as Linker

#### Part 1: Functionalization with AEAPMS

**1|** Weigh 100 mg of $TiO_2$ nanotubes ($TiO_2$-NTs) into a 100 mL round-bottom flask.

**2|** Add 30 mL of dry toluene to the flask with nanotubes and sonicate in the knot of a sonic bath (e.g. Elma, Elmasonic P) for 20 minutes at 37 kHz and power set to 320 W to ensure a good dispersion.
▲ **CRITICAL STEP** Dry toluene must be used to avoid having too much water in the reaction mixture, which can cause cross-linking of AEAPMS.

**3|** Add a magnet to the flask and set the flask on a magnetic stirrer-heater. Heat the dispersion on an oil bath to 60 °C at constant stirring.

**4|** Pour 30 mL of dry toluene into a fresh 50 mL round bottom flask, then add 840 µL of 3-(2-aminoethylamino)propyltrimethoxysilane (AEAPMS). Vortex/shake the solution for 30 seconds for thorough mixing.
▲ **CRITICAL STEP** Again, dry toluene must be used to avoid traces of water.

**5|** In 30 minutes time, dropwise add the solution of AEAPMS diluted in dry toluene to the dispersion of $TiO_2$-NTs in dry toluene.

**6|** The reaction mixture is kept at 60 °C for one week.

**7|** Afterwards, the reaction mixture is cooled down to room temperature by turning the heater off.

**8|** Centrifuge the reaction mixture in a 100 mL glass centrifuge tube for 5–10 minutes at 2500 rpm.

**9|** Remove/decant the liquid phase.

**10|** Rinse the solid material with 30 mL of toluene and centrifuge for 5–10 minutes at 2500 rpm. After centrifugation decant/remove the liquid phase. Repeat 3 times.

**11|** Repeat step **10** with hexane instead of toluene. Rinse 3 times as well.

**12|** Place the glass centrifuge tube in the oven and dry the product at 80 °C overnight (12 h).

**13|** Transfer the glass centrifuge tube into a vacuum drier and dry the product at 80 °C, 150 mbar for 2 h.

**14|** Store the functionalized $TiO_2$ nanotubes (f-$TiO_2$-NTs) in plastic boxes with caps or in glass vials. Use parafilm to additionally seal the content to avoid possible accidents.

➤ **QUALITY CHECK** The quality of functionalization is verified with FTIR and Zeta potential measurements of both the native and the functionalized material.

**Part 2: Labelling with Alexa Fluor 488 SDP ester**

This labelling protocol is designed for labelling of AEAPMS-functionalized $TiO_2$ nanotubes with Alexa Fluor 488 SDP ester hydrophilic dye. If choosing a different linker and/or dye, several changes should be made to the protocol. Most importantly, the pH of the reaction solution should assist the chemical reaction between the linker and dye, while at the same time ensure a stable nanoparticle suspension.

Part 2a: Buffer preparation

This recipe is for a 100-times diluted bicarbonate buffer (100x dcb) with pH 10 and osmolarity 5 mOsm/L which was found to work well for labelling of our functionalized $TiO_2$ nanotubes with negatively charged SDP ester dyes. The range of optimal pH values depends on the zeta potential of the functionalized $TiO_2$ nanotubes and the functional group on the dye.

**15|** Weigh 2.65 g of $NaHCO_3$ and 2.10 g of $Na_2CO_3$ to a 500 mL Florence flask.

**16|** Add 500 mL of purified water "Type 1" (e.g. MilliQ water) and a magnet to the 500 mL Florence flask.

**17|** Set the 500 mL Florence flask to magnetic stirring for 10 min at 1000 rpm. The Florence flask now contains sodium bicarbonate buffer with pH ≈ 10 and osmolarity 0.5 Osm/L.

**18|** Measure the pH of the buffer with a pH electrode. If values less than 9.8 are obtained, the buffer should be prepared again with greater care or using an adjusted ratio of salts in accordance with the pH of the "Type 1" water.

**▲ CRITICAL STEP** At pH lower than 9.5 the $TiO_2$ nanotubes_will be more prone to agglomeration and the majority of them will be poorly dispersed (the critical pH can be deduced from the Zeta potential measurement of f-$TiO_2$-NTs). During labelling we wish to have as much surface of nanotubes exposed to the dye as possible.

**19|** Dilute some of the buffer 100 times by addition of 1 mL of buffer and 99 mL of purified (e.g. MilliQ) water to a 100 mL Florence flask. You have obtained 100-times diluted sodium bicarbonate buffer (100x dcb) with osmolarity 5 mOsm/L. Save the rest of the non-diluted bicarbonate buffer for later use.

**20|** Measure the pH of 100x dcb with pH-meter. The measured pH should be around 10 – if not, prepare the dilution again.

Part 2b: Nanomaterial Scaling

**21|** Weigh 6 mg of f-$TiO_2$-NTs using a high-precision scale, directly to an 8 mL glass vial.

**22|** Add 6 mL of 100x dcb with pH 10 (prepared in steps 1-6) to the 8 mL glass vial to obtain a nanotube concentration of 1 mg/mL.

**▲ CRITICAL STEP** The initial solution of 0.1 M sodium bicarbonate buffer with pH ≈ 10 has to be at least 100-times diluted because of the charge screening between surface of f-$TiO_2$-NTs and the fluorescent dye. If the osmolarity of the buffer is too high, electrostatic attraction between the dye and the linker becomes negligible at distances larger than 1–10 nm. This results in unsuccessful labelling of f-$TiO_2$-NTs as the probability of the reaction between the SDP esters and linker diminishes and a vast majority of the dyes react with water molecules instead of binding to the linker. Also, the nanoparticle concentration must not exceed 1 mg/mL. Larger concentration of nanoparticles will result in over-packing of nanoparticles and thus steric confinement of nanoparticles. This results in a significant reduction of exposed nanoparticle surface and, as a consequence, greatly reduces the efficiency of labelling.

**23|** Sonicate the 8 mL glass vial with a dispersion of f-$TiO_2$-NTs on a sonication bath (e.g. Branson 2510) for 30 s for initial dispersal of nanotubes.

Part 2c: Preparing the Tip Sonicator

**24|** Unscrew the microtip from the tip sonicator (e.g. MISONIX Ultrasound liquid Processor with 419 MicrotipTM) and polish it with fine sand paper on an even surface until smooth.

**▲ CRITICAL STEP** Be careful to thoroughly wash the microtip afterwards to remove all debris from the polishing.

**25|** Screw the microtip back onto the sonicator.

**26|** Add 6 mL of 96% ethanol (EtOH) to a 12 mL plastic centrifuge tube (Falcon tube) and mount it to the stand inside the sonicator box.

**27|** Adjust the 12 mL plastic centrifuge tube so that approximately 4 cm of the microtip is submerged in EtOH.

**28|** Sonicate on ice bath for 1 min run-time with a 5 s on, 5 s off regime (2 min total time) with a power of 20-30 W (on the MISONIX Ultrasound liquid Processor with 419 MicrotipTM this corresponds to an amplitude value of 70) .

**29|** Remove the 12 mL plastic centrifuge tube with EtOH and wipe the tip with a fresh paper towel.

**30|** Add 6 mL of 100x dcb with pH ≈ 10 to another 12 mL plastic centrifuge tube and mount it to the stand inside the sonicator box.

**31|** Repeat steps **27-29** using the 100x dcb instead of EtOH for additional washing of the tip. This time, do not wipe the tip after sonication.

Part 2d: Dispersing the Nanomaterial

**32|** Mount the 8 mL glass vial with the dispersion of f-$TiO_2$-NTs to the stand.

**33|** Adjust the 8 mL glass vial so that the tip is approximately 1 cm from the bottom of the vial and is not touching the glass vial walls. Tightly seal the opening of the vial around the sonicator tip with parafilm. Close the sonicator door, then close and run the fume hood.

**▲ CRITICAL STEP** Special care must be taken to prevent the sonicator tip from touching the glass vial - sparks may be produced at the contact of microtip with glass, the vibrations on the glass vial may cause the vial to slip from iron stage clamp and fall into the ice bath, which will result in loss of sample. In the most severe case, the vibrations could cause the glass vial to break.

**34|** Sonicate on ice bath for 15 min run time with 5 s on, 5 s off regime (30 min total time). The power should again be 20–30 W (amplitude 70).

Part 2e: Labelling the Nanomaterial

**35|** Dilute 1 mg of fresh Alexa Fluor 488 SDP ester in 100 μL of anhydrous DMSO to obtain concentration of 12.1 mM.

▲ **CRITICAL STEP** It is very important to have fresh dye or dye that has been stored in powder form at –20 °C or lower, and has not been in contact with humid air or any liquid containing traces of water. Ester groups are very reactive to water, which consequently leads to loss of ester binding groups, necessary for attachment of the dye to the linker. Effectiveness of labelling is thus severely lowered. The solution of dye should be made right before adding it to freshly sonicated functionalized nanomaterial.

**36|** Add 19.8 μL of 12.1 mM Alexa Fluor 488 to the 8 mL glass vial containing freshly sonicated nanotubes. This corresponds to 40 nmol of dye per 1 mg of nanoparticles, which we estimated to be a few-fold more dye than the number of designated linker targets.

**37|** Vortex the 8 mL glass vial gently for 15 seconds.

**38|** Repeat steps **33** and **34**. Change the run time to 30 min (60 min total). Before running the sonication, cover everything with aluminum foil to prevent photo-bleaching of the dye.

**39|** Remove the 8 mL glass vial from the iron stage clamp, add a small magnet and close the vial with a cap. Use parafilm to seal the cap tightly.

**40|** Set the 8 mL glass vial to a magnetic mixer, set to 260 rpm. Secure the vial firmly in place so it does not tip over, and wrap everything tightly in aluminum foil. Leave the vial on the mixer overnight (12–14 h).

**41|** After 12–14 h remove the 8 mL glass vial from the magnetic mixer. Remove the magnet with a magnet picker without touching the sample.

**42|** Close the vial and seal it with parafilm. Wrap the vial in aluminum foil and keep in the refrigerator (4 °C) until free dye removal (part 3) is performed.

**Part 3: Free dye removal with a centrifugal filter device**

This protocol is designed for cleaning 2 mL of suspended labelled nanoparticles using a 4 mL Amicon Ultra 100K centrifugal filter device (CFD) with a 100 kDa membrane. Up to 4 mL can easily be filtered using the same CFD.

▲ **CRITICAL STEP** If using a different filter device and/or different volume of nanoparticles, appropriate adjustments must be made to centrifugation times and speeds, medium volumes, etc. Also, the pore size of the membrane in the centrifugal filter device must be larger than the dye and smaller than the size of nanoparticles (see manufacturer’s guidelines).

▲ **CRITICAL STEP** If different dyes and nanoparticles are used, the cleaning medium might need to be swapped for a more appropriate one (the nanoparticles must be in a stable dispersion and the dye must be soluble in the medium). Note that the last two filtrations should use the final, storage medium.

Part 3a: Free dye removal

**43|** Prepare the centrifugal filter device according to the manufacturer's guidelines – for instance, wet the membrane by centrifuging 1 mL of deionized water through it. Afterwards, decant the water and dry the filtrate compartment with a lab-grade paper tissue (this prevents false readouts of dye concentration in the filtrates due to dilution).

**44|** Place 2 mL (depends on the filtration device you are using) of your labelled nanoparticles into the upper part of the device and close the cap. Place the device inside the centrifuge and balance it with a centrifugal tube, filled with water.

**45|** Centrifuge for 15 minutes at 1000 g (or according to the manufacturer's guidelines).

**46|** When the centrifugation is complete, take the centrifugal filter device out of the centrifuge and transfer the filtrate (liquid that accumulated below the membrane) into an Eppendorf for further analysis and close it with parafilm (to prevent evaporation). You should also wrap your samples in aluminum foil to prevent photo-bleaching.

**47|** Fill the filtrate compartment with a medium, consisting of ethanol and bicarbonate buffer in 70:30 mass ratio and the same osmolarity as 100x dcb ("70% EtOH 100x dcb", recipe: 31.69 g of 96% EtOH is weighed (or 39.37 mL is measured), then 0.5 mL non-diluted bicarbonate buffer and 11.27 g (11.27 mL) of $H_2O$ are added). Use enough of this mixture so that when you put the membrane back into the filter device, the level of the medium in the filtrate compartment reaches up to the 2 mL mark – when you will resuspend the nanoparticles in the retentate compartment with 2 mL of medium, the height of both media will be about the same. This step ensures better cleaning of the membrane with the ultrasound bath in step 49. The medium and its level should be the same in both compartments to avoid changing composition and/or volume in the retentate compartment.

**48|** Resuspend the sample in the upper part of the device (the retentate) with 2 mL of 70% EtOH 100x dcb (same medium as step **47**) and mix thoroughly with pipette.

**49|** Close the filter device and sonicate on sonication bath (e.g. Branson 2510) for 10 seconds to diminish the amount of nanoparticles stuck to the CFD membrane and to break up the nanoparticle aggregates.

**50|** Discard the content of the filtrate compartment.

**51|** Wash the filtrate compartment of the device using EtOH and wipe it with a lab-grade paper tissue. This prevents false readouts of dye concentration in the filtrates.

**52|** Repeat steps **45-51** once again with 70% EtOH 100x dcb. Then, repeat the steps two more times with 100x dcb to reduce the amount of ethanol in the final sample. Then, repeat steps **45 – 47** one final time.

Part 3b: Collecting the final, cleaned sample

**53|** Fill the filtrate compartment with the desired final medium (we usually use 100x dcb). Again, use enough of it so that when you put the membrane into the CFD, it is submerged up to the 2 mL mark.

**54|** Resuspend the retentate in half of the starting volume (in our case, 1 mL) of the desired final medium and mix thoroughly with a pipette.

**55|** Close the filter device and sonicate on sonication bath (e.g. Branson 2510) for 10 seconds to diminish the amount of nanoparticles stuck to the CFD membrane and to break up the nanoparticle aggregates.

**56|** Discard the content of the filtrate compartment.

**57|** Transfer the retentate into a 2 mL Eppendorf (or glass vial).

**58|** Repeat steps **54-57** once more so an appropriate concentration of nanoparticles is achieved (transfer the sample into the same Eppendorf as in step **57** so the final concentration of nanoparticles is the same as in the beginning).

**59|** Check that all the saved filtrates and retentates are properly labelled, closed with parafilm and wrapped in aluminum foil to prevent photo-bleaching.

➤ **QUALITY CHECK** The success of cleaning is verified with calculation of the free dye concentration in the filtrates via fluorometric measurements or from FCS measurements. For reference, the fluorescence signal from the non-cleaned and fully cleaned sample are measured as well.

**60|** If needed, repeat steps **45-47**, **53-59** prior to measurement with the labelled nanoparticles to eliminate any dye which might have desorbed from the nanoparticles. In our case the cleaned sample was stable for up to a few weeks. After this time, the amount of free dye interfered with the measurements unless it was cleaned with the centrifugal filter device prior to the measurement. Also repeat the steps **45-47**, **53-59** twice if you need to exchange storage medium for another medium.
